# Global analysis of gene expression dynamics identifies factors required for accelerated mRNA degradation

**DOI:** 10.1101/254920

**Authors:** Darach Miller, Nathan Brandt, David Gresham

## Abstract

Cellular responses to changing environments frequently involve rapid reprogramming of the transcriptome. Regulated changes in mRNA degradation rates can accelerate reprogramming by clearing or stabilizing extant transcripts. Here, we measured mRNA stability using 4-thiouracil labeling in the budding yeast *Saccharomyces cerevisiae* during a nitrogen upshift and found that 78 mRNAs are subject to destabilization. These transcripts include Nitrogen Catabolite Repression (NCR) and carbon metabolism mRNAs, suggesting that mRNA destabilization is a mechanism for targeted reprogramming. To explore the molecular basis of destabilization we implemented a SortSeq approach to screen using the pooled deletion collection library for *trans* factors that mediate rapid *GAP1* mRNA repression. We combined low-input multiplexed Barcode sequencing with branched-DNA single-molecule mRNA FISH and Fluorescence-activated cell sorting (<u>BFF</u>) to identify that the Lsm1-7p/Pat1p complex and general mRNA decay machinery are important for *GAP1* mRNA clearance. We also find that the decapping modulator *SCD6,* translation factor eIF4G2, and the 5’ UTR of *GAP1* are important for this repression, suggesting that translational control may impact the post-transcriptional fate of mRNAs in response to environmental changes.

## Introduction

Regulated changes in mRNA abundance are a primary cellular response to external stimuli. Both the rate of synthesis and the rate of degradation determine the steady-state abundance of a particular mRNA and the kinetics with which abundance changes occur (*Hargrove and Schmidt, 1989; Pérez-Ortín et al., 2013*). Changes in mRNA degradation rates fulfill an important mechanistic role in diverse systems, including development (*Alonso, 2012; West et al., 2017*) and disease (*Aghib et al., 1990*). In budding yeast, the rate of mRNA degradation is affected by environmental stresses (*Canadell et al., 2015*), cellular growth rate (*García-Martínez et al., 2016*), as well as by improvements in nutrient conditions (*Scheffler et al., 1998*).

Environmental shifts trigger rapid reprogramming of the budding yeast transcriptome in response to stresses and nutritional changes (*Gasch et al., 2000; Conway et al., 2012*). mRNA degradation rate changes have been shown to play a role in responses to heat-shock, osmotic stress, pH increases, and oxidative stress (*Castells-Roca et al., 2011; Romero-Santacreu et al., 2009; Canadell et al., 2015; Molina-Navarro et al., 2008*). In response to these diverse stresses destabilization of mRNAs encoding ribosomal-biogenesis gene products, and stress-induced mRNA occurs (*Canadell et al., 2015*). Simultaneous increases in both synthesis and degradation rates of some mRNAs may serve to speed the return to a steady-state following a transient pulse of regulation (*Shalem et al., 2008*). Addition of glucose to carbon-limited cells results in both stabilization of ribosomal protein mRNAs (*Yin et al., 2003*) and destabilization of gluconeogenic transcripts (*de la Cruz et al., 2002; Mercado et al., 1994*). Destabilization of transcripts can have a delayed effect on reducing protein levels compared to up-regulated genes (*Lee et al., 2011*). This suggests that accelerated mRNA degradation may serve additional purposes. For example, clearance of specific mRNAs could increase nucleotide pools (*Kresnowati et al., 2006*) or facilitate reallocation of translational capacity (*Kief and Warner, 1981; Giordano et al., 2016; Shachrai et al., 2010*).

Yeast cells metabolize a wide variety of nitrogen sources, but preferentially assimilate and metabolize specific nitrogen compounds. Transcriptional regulation, known as “nitrogen catabolite repression” (NCR) (*Magasanik and Kaiser, 2002*), controls the expression of mRNAs encoding transporters, metabolic enzymes, and regulatory factors required for utilization of alternative nitrogen sources. NCR-regulated transcripts are expressed in the absence of a readily metabolized (preferred) nitrogen sources or in the presence of growth-limiting concentrations (in the low μM range) of any nitrogen source (*Godard et al., 2007; Airoldi et al., 2016*). Regulation of NCR targets is mediated by two activating GATA transcription factors, Gln3p and Gat1p, and two repressing GATA factors, Dal80p and Gzf3p. *GAT1, GZF3,* and *DAL80* promoters contain GATAA motifs, and thus transcriptional regulation of NCR targets entails self-regulatory and cross-regulatory loops. When supplied with a preferred nitrogen source such as glutamine, the NCR-activating transcription factors Gat1p and Gln3p are excluded from the nucleus by TORC1-dependent and-independent mechanisms (*Beck and Hall, 1999; Tate and Cooper, 2013; Tate et al., 2017*) and NCR transcripts are strongly repressed. The activity of some NCR gene products is also controlled by post-translational mechanisms (*Cooper and Sumrada, 1983*) such as the General Amino-acid Permease (Gap1p) which is rapidly inactivated upon a nitrogen upshift via ubiquitination (*Stanbrough and Magasanik, 1995; Risinger et al., 2006; Merhi and Andre, 2012*). Recently, we have identified an additional level of regulation of NCR transcripts: cells growing in NCR de-repressing conditions accelerate the degradation of *GAP1* mRNA upon addition of glutamine (*Airoldi et al., 2016*). Thus, mRNA degradation rate regulation may be an additional mechanism for clearing NCR-regulated transcripts upon improvements in environmental nitrogen availability.

Multiple pathways mediate the degradation of mRNAs. The main pathway of mRNA degradation occurs by deadenylation and decapping prior to 5’ to 3’ exonucleolytic degradation by Xrn1p; however, transcripts are also degraded 3’ to 5’ via the exosome, or via activation of co-translational quality control mechanisms (*Parker, 2012*). Deadenylation of mRNAs by the Ccr4-Not complex allows the mRNA to be bound at the 3’ end by the Lsm1-7p/Pat1p complex, a heptameric ring comprising the SM-like proteins Lsm2-7p and the cytoplasmic-specific Lsm1p (*Tharun et al., 2000; Sharif and Conti, 2013*), which then recruits factors for decapping by Dcp2p. Recruitment of the decapping enzyme (*Coller and Parker, 2004*) is the rate-limiting step for canonical 5’-3’ degradation. Therefore Lsm1-7p, Pat1p, and associated factors play a key role (*Nissan et al., 2010*).

Regulation of mRNA degradation pathways can alter the stability of specific mRNAs. For example, the RNA-binding protein (RBP) Puf3p recognizes a cis-element in 3’ UTRs (*Olivas and Parker, 2000*) and affects mRNA degradation rates depending on Puf3p phosphorylation status (*Lee and Tu, 2015*). In addition to cis-elements within the transcirpt, promoters have been shown to mark certain RNA-protein (RNP) complexes to specify their post-transcriptional regulation (*Mercado et al., 1994; Haimovich et al., 2013; Trcek et al., 2011; Braun et al., 2015*). These mechanisms may be controlled by a variety of different signalling pathways including Snf1 (*Young et al., 2012; Braun et al., 2014*), PKA (*Ramachandran et al., 2011*), Phk1/2 (*Luo et al., 2011*), and TORC1 (*Talarek et al., 2010*). Thus, regulated changes in mRNA degradation rates entails numerous mechanisms that collectively tune stability of mRNAs in response to the activity of signalling pathways.

Here, we studied the global regulation of mRNA degradation rates upon improvment in environmental nitrogen using 4-thiouracil (4tU) label-chase and RNAseq. We found that a set of 78 mRNAs are subject to accelerated mRNA degradation, including many NCR transcripts as well as mRNAs encoding components of carbon metabolism. To identify the mechanism underlying accelerated mRNA degradation we designed a high-throughput genetic screen using <u>B</u>arcode-sequencing of a pooled library which was fractionated using Fluorescence-activated cell sorting of single molecule mRNA <u>F</u>ISH signal (BFF). We screened the barcoded yeast deletion collection to test the efect of each gene deletion on the abundance of *GAP1* mRNA in NCR de-repressing conditions and its clearance following the addition of glutamine. We find that the Lsm1-7p/Pat1p complex and decapping modifiers affect both *GAP1* mRNA steady-state expression and its accelerated degradation. This work expands our understanding of mRNA stability regulation in remodeling the transcriptome during a relief from growth-limitation and demonstrates a generalizable approach to the study of genetic determinants of mRNA dynamics.

## Results

### Transcriptional reprogramming precedes physiological remodeling

Cellular responses to environmental signals entail coordinated changes in both gene expression and cellular physiology. Previously, we studied the steady-state and dynamic responses of *Saccharomyces cerevisiae* (budding yeast) to environmental nitrogen (*Airoldi et al., 2016*), and found that the transcriptome is rapidly reprogrammed following a single pulsed addition of glutamine to nitrogen-limited cells in either a chemostat or batch culture. To study physiological changes in response to a nitrogen upshift, we measured growth rates of a population of cells. A prototrophic haploid lab strain (FY4, isogenic to S288c) grows with a 4.5 hour doubling time in batch culture in minimal media containing proline as a sole nitrogen source (*Figure 1a*). Upon addition of 400μM glutamine the cells undergo a 2-hour lag period during which no change in population growth rate is detected, but the average cell size continuously increases (∼21% increase in mean volume *Figure 1b*). Following the lag, the population adopts a 2.1 hour doubling time. By contrast, global gene expression changes are detected within three minutes of the upshift (*Airoldi et al., 2016*). Thus, transcriptome remodeling precedes physiological remodeling in response to a nitrogen upshift.

**Figure 1.**
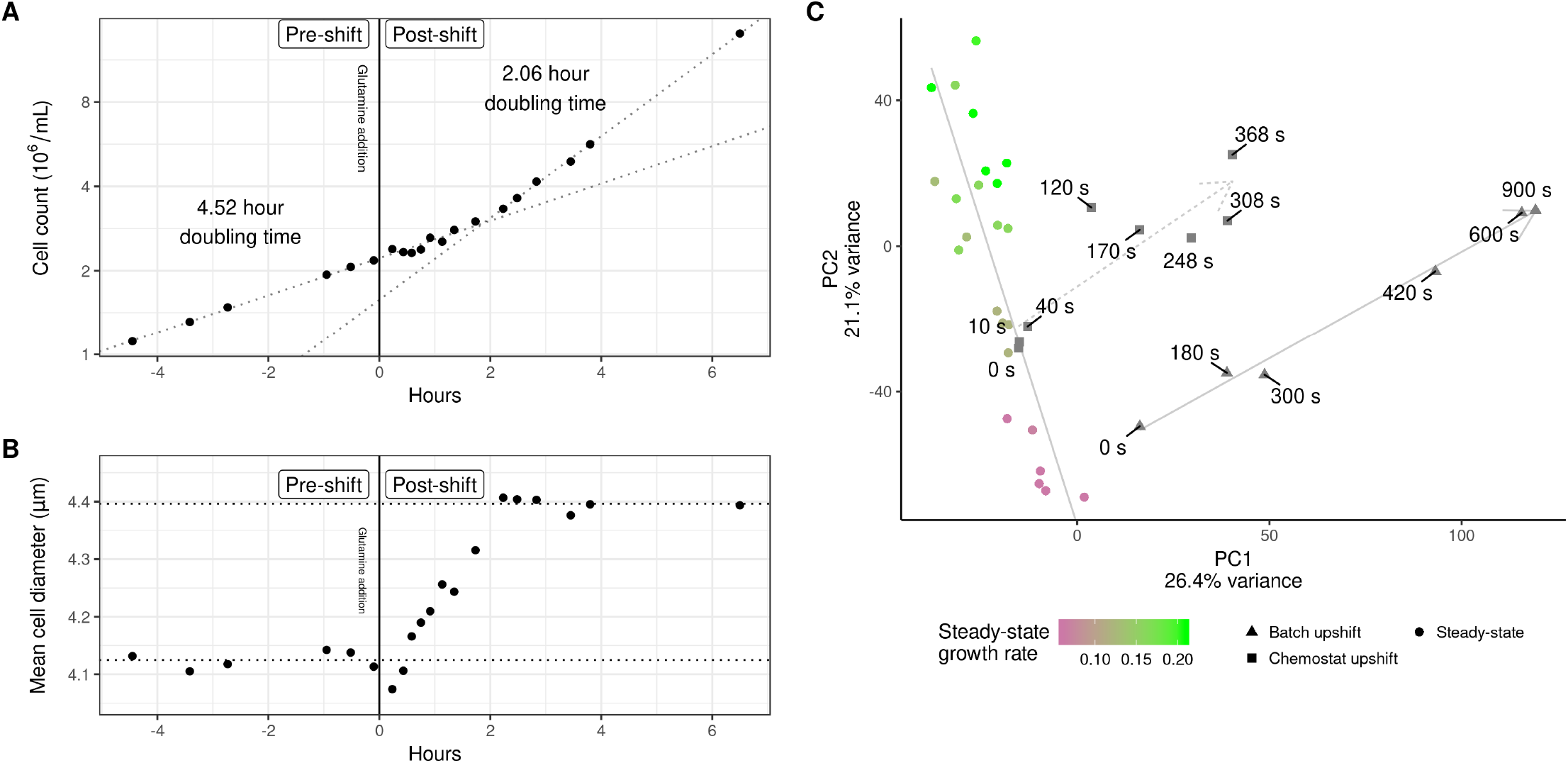
Dynamics of physiological and transcriptome remodeling during a nitrogen upshift. **a)** 400μM glutamine was added to a culture of yeast cells growing in minimal media containing 800μM proline as a sole nitrogen source. Measurements of culture density across the upshift are plotted. Dotted lines denote linear regression of the natural log of cell density against time before the upshift and after the 2 hour lag. **b)** Average cell size. Dotted lines denote the mean cell diameter before the upshift and after the 2 hour lag. **c)** PCA analysis of global mRNA expression in steady-state chemostats and following an upshift (Airoldi et al., 2016). Steady-state nitrogen-limited chemostat cultures maintained at different growth rates (colored circles) primarily vary along principal component 2. Expression following a nitrogen-upshift in either a chemostat (squares) or batch culture (triangles) show similar trajectories and primarily vary along principal component 1. Grey lines depict the major trajectory of variation for the steady-state and upshift experiments. **Figure 1–Figure supplement 1.** Long-term transcriptome dynamics upon nitrogen upshift. **Figure 1–Figure supplement 2.** Gene set enrichment analysis of loadings on principal components one and two.

To evaluate concordance in transcriptome remodeling between chemostat and batch nitrogen upshifts, and the extent to which they reflect changes in gene expression observed during systematic steady-state changes in growth rates using chemostats, we performed principal component analysis of global gene expression (*Figure 1c*). The first two principal components, which account for almost half of the total variation, clearly separate steady-state and nitrogen upshift cultures. Systematic changes in growth rate primarily results in separation of gene expression states along the second principal component, whereas upshift experiments vary along the first principal component. This suggests that although a nitrogen upshift results in a gene expression state reflecting increased cell growth rates (*Airoldi et al., 2016*), the transcriptome is remodeled through a distinct state. In upshift experiments in chemostats, the gene expression trajectory ultimately returns to the initial steady-state condition as excess nitrogen is depleted by consumption and dilution (*Figure 1-Figure Supplement 1*).

To investigate the functional basis of gene expression programs in the upshift and steady-state conditions, we computed the correlation of each transcript with the loadings on these first two principal components and performed gene-set enrichment analysis (*Figure 1-Figure Supplement 2*). Component 1 is positively correlated with functions like mRNA processing, transcription from RNA polymerases (I,II,and III), and chromatin organization, and negatively correlated with cytoskeleton organization, vesicle organization, membrane fusion, and cellular respiration. Both steady-state and upshift gene expression trajectories increase with principal component 2, but they diverge along principal component 1. Components 1 and 2 are strongly enriched for terms including ribosome biogenesis, nucleolus, and tRNA processing, and negatively correlated with vacuole, cell cortex, and carbohydrate metabolism terms. Together, this analysis suggests that both upshift and increased steady-state growth rates share upregulation of ribosome-associated components, but the reprogramming preceeding the upshift in growth reflects an immediate focus on gene expression machinery instead of structural cellular components. Importantly, dynamic reprogramming is similar in both the chemostat and batch upshift (*Figure 1c*). As batch cultures are a technically simpler experimental system, we performed all subsequent experiments using batch culture nitrogen upshifts.

### Global analysis of mRNA stability changes during the nitrogen upshift

Previously, we found that *GAP1* and *DIP5* mRNAs are destabilized in response to a nitrogen upshift (*Airoldi et al., 2016*). We sought to determine if mRNA destabilization is specific to NCR transporter mRNAs by measuring global mRNA stability across the upshift using 4-thiouracil (4tU) labeling and RNA-seq (*Neymotin et al., 2014; Munchel et al., 2011*). As 4tU labeling requires nucleotide transport, which may be altered upon a nitrogen-upshift (*Hein et al., 1995*), we designed experiments such that following complete 4tU labeling and metabolism to nucleotides the chase was initiated prior to addition of glutamine or water (mock). We normalized data using *in vitro* synthesized thiolated spike-ins by fitting a log-linear model to spike-in counts across time (*Figure 2-Figure Supplement 7*), which reduced noise and increased our power to detect stability changes ( *Figure 2-Figure Supplement 8, Figure 2-Figure Supplement 9, Figure 2-Figure Supplement 10*). Data and models for each transcript can be visualized in browser using a Shiny appplication (see http://shiny.bio.nyu.edu/users/dhm267/ or *Appendix 1*).

We modeled the log-transformed normalized signal for each mRNA using linear regression (*Figure 2-Figure Supplement 12*). Of 4,859 mRNAs measured we identified 94 that increased in degradation rate and 38 that decreased (FDR < 0.01, using *Storey and Tibshirani (2003*)). We generated a model of nucleotide labeling kinetics to assess the effect of an incomplete label chase on our experimental design ( *Figure 2-Figure Supplement 7*), and found that complete transcriptional inhibition alone could theoretically result in a 17% increase in the apparent degradation rate. In order to eliminate the possibility that rapid synthesis changes could affect our estimates, we only considered destabilization of at least a doubling (100% increase) of apparent degradation rates between pre-upshift and post-upshift. This conservative cutoff left 78 mRNA that are significantly destabilized upon a nitrogen upshift.

The vast majority of transcripts (4,781 of 4,859) do not show individual evidence for stability changes upon addition of glutamine (e.g. *HTA1, Figure 2a*). The median pre-upshift half-life is 6.89 minutes and the median half-life following the upshift is 6.4 minutes (*Table 1*) suggesting that there is not a global change in mRNA stability. Global stability estimates are considerably lower than previous estimates in rich medium (*Munchel et al., 2011; Neymotin et al., 2014; Miller et al., 2011*), which may reflect the different nutrient conditions used in our study. The 78 transcripts significantly destabilized upon the glutamine-upshift include mRNAs encoding NCR transporters *GAP1, DAL5,* and *MEP2* (blue label, *Figure 2a*), the pyruvate metabolism enzymes *PYK2* and *PYC1* (orange label), and trehalose synthase subunits *TPS1* and *TPS2* (yellow label). Destabilized mRNA tend to be more stable before the upshift (*Figure 2b*), (median half-life of 9.46 minutes) and exhibit a median 3.06-fold increase in degradation rates (median half-life of 3.02 minutes following the upshift).

**Figure 2.**
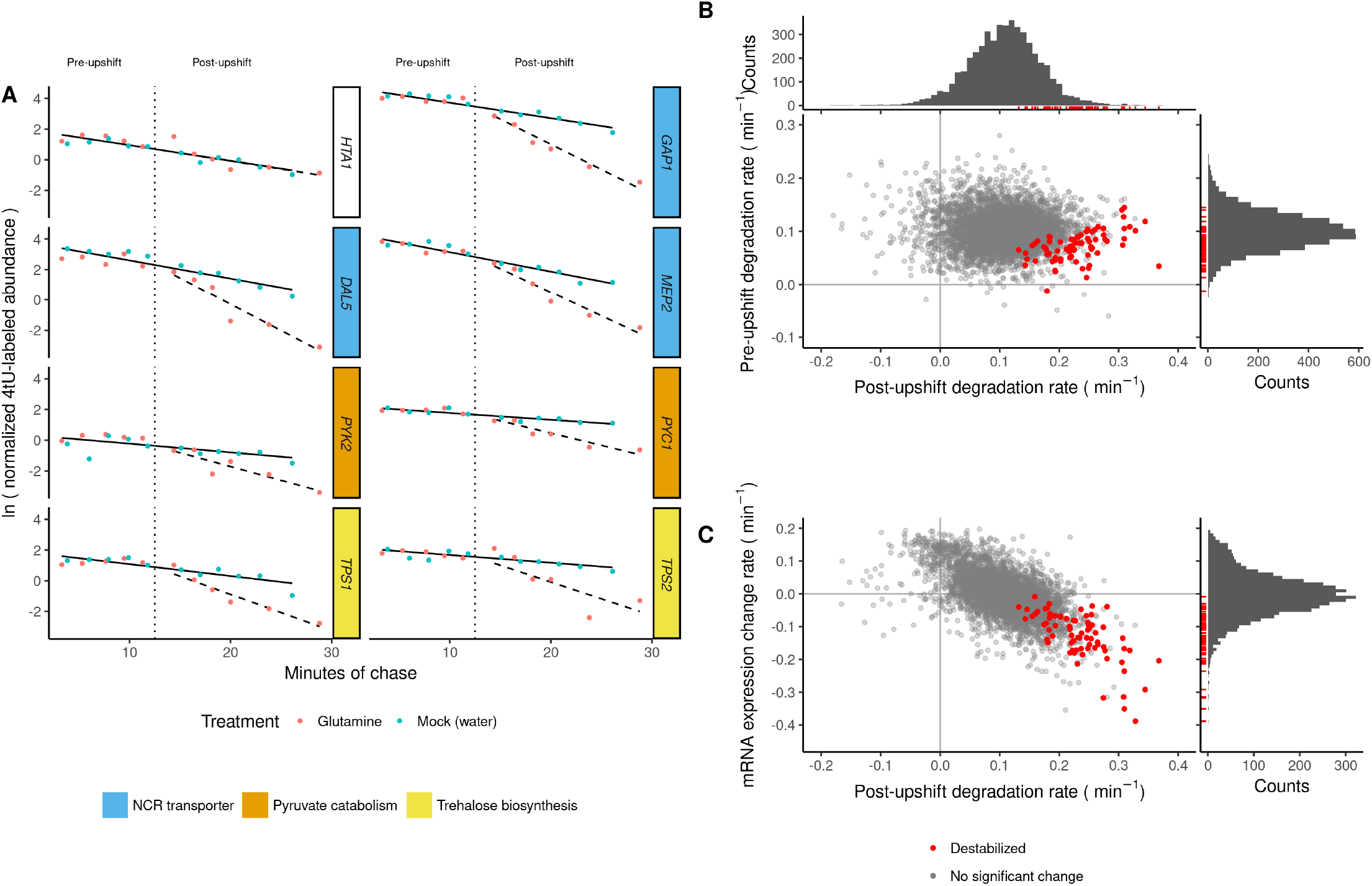
Global mRNA stability change following a nitrogen upshift. **a)** 4tU-labeled mRNA from each gene was measured over time, before and after the addition (vertical dotted line) of glutamine (nitrogen-upshift) or water (mock). Linear regression models were fit to the data with a rate before the upshift (solid line) and a rate after glutamine addition (dashed line). *HTA1* is not significantly destabilized, whereas mRNAs encoding NCR-regulated transporters or pyruvate and trehalose metabolism enzymes are significantly destabilized. **b)** Comparison between the pre-upshift mRNA degradation rate (y-axis) and the post-upshift mRNA degradation rate (x-axis). Positive values result from noise on the slope estimate. **c)** Comparison between changes in mRNA expression following upshift (Airoldi et al. 2016) (y-axis) and the post-upshift degradation rate (x-axis). Both plots share the same x-axis. Transcripts that are significantly destabilized are colored red, and shown with red rug-marks in the marginal histograms. **Figure 2-Figure supplement 1.** Enrichment of Hrp1p motif in 5’ UTRs of destabilized transcripts. **Figure 2-Figure supplement 2.** Comparison between rates of mRNA abundance change (*Airoldi et al., 2016*) and stability measured in this study. **Figure 2-Figure supplement 3.** Many of the destabilized mRNA are members of the ESR-up regulon (*Gasch et al., 2000*). **Figure 2-Figure supplement 4.** Abundance changes versus stability, plotted for only the destabilized transcripts. **Figure 2-Figure supplement 5.** Six examples of individual mRNA whose regulation is more complex than a homo-directional destabilization and synthesis repression. **Figure 2-Figure supplement 6.** The destabilized set is longer and has a higher frequency of optimal codons than the rest of the transcriptome. **Figure 2-Figure supplement 7.** Supplementary file with experimental rationale, details, and protocol for the label-chase experiment. **Figure 2-Figure supplement 8.** Raw counts of labeled mRNA quantified by RNAseq in label-chase experiment. **Figure 2-Figure supplement 9.** Filtered label-chase RNAseq data for modeling, normalized directly within sample. **Figure 2-Figure supplement 10.** Filtered label-chase RNAseq data for modeling, normalized by modeling across samples. **Figure 2-Figure supplement 11.** Degradation rate modeling results, from data normalized within samples. **Figure 2-Figure supplement 12.** Degradation rate modeling results, from data normalized across samples. **Figure 2-Figure supplement 13.** Enriched GO and KEGG terms within the set of mRNA destabilized upon a nitrogen upshift, across sample normalization.

**Table 1.** Summary of mRNA stability, median values

|  | Pre-shift |  | Post-shift |  | Change in specific rate (min<sup>-1</sup>) | Fold-change specific rate |
| --- | --- | --- | --- | --- | --- | --- |
|  | specific rate (min<sup>-1</sup>) | half-life (min) | specific rate (min<sup>-1</sup>) | half-life (min) |  |  |
| All transcripts | 0.100 | 6.92 | 0.110 | 6.32 | 0.00865 | 1.08 |
| Destabilized (n=78) | 0.0732 | 9.46 | 0.229 | 3.02 | 0.158 | 3.06 |
| Destabilized (n=4781) | 0.101 | 6.89 | 0.108 | 6.40 | 0.00728 | 1.07 |

We tested for functional enrichment among the set of 78 destabilized mRNAs and found that they are strongly enriched for NCR transcripts (16 of 78, p < 10^−11^). Almost half of the destabilized transcripts are annotated as “ESR-up” genes (*Figure 2-Figure Supplement 3*), on the basis of their rapid induction during the environmental stress response (*Gasch et al., 2000*). These 78 destabilized mRNA are enriched (FDR < 0.05) for GO terms and KEGG pathways (*Figure 2-Figure Supplement 13*) including glycolysis/gluconeogenesis (6 genes), carbohydrate metabolic process (24), trehalose-phosphatase activity (3), pyruvate metabolic process (6), and secondary active transmembrane transport (8, a subset of the enriched 11 ion transmembrane transport genes). Thus destabilized mRNA upon a nitrogen upshift regulates, but is not restricted to, NCR-regulated mRNA and reflects broader metabolic changes in the cell.

To investigate the extent to which mRNA stability changes contribute to transcriptome reprogramming, we compared degradation rates to abundance changes following the upshift (*Airoldi et al. (2016), Figure 2c*). Changes in mRNA degradation rates and expression change rates are anti-correlated (Pearson’s *r* = -0.598, p-value < 10^−15^, *Figure 2-Figure Supplement 2*), consistent with stability changes impacting gene expression dynamics. However, they are not entirely co-incident, as some destabilized transcripts do not exhibit decreases in abundance (red points in *Figure2 c, Figure 2-Figure Supplement4,* and *Figure 2-Figure Supplement5*). This analysis shows that increases in degradation rates play a significant role in the rapid reprogramming of the transcriptome upon a glutamine upshift, but that in some cases cases they are counteracted by increases in mRNA synthesis rates (*Shalem et al., 2008; Canadell et al., 2015*).

Functional coordination of mRNA stability changes suggests a possible role for *cis*-element regulation. We analyzed UTRs and coding sequence for enrichment of new motifs or known RNA binding protein (RBP) motifs. 3’ UTRs of destabilized transcripts are depleted of Puf3p binding sites, and we found no enriched sequence motif in the 3’ UTRs. 5’ UTRs are enriched for a GGGG motif, which may be explained by convergence between mRNA stability changes and transcriptional control by Msn2/4 on the ESR “up” genes (*Figure 2-Figure Supplement 3, Gasch et al. (2000); Canadell et al. (2015*)). 5’ UTRs are also enriched for binding motifs reported for Hrp1p (*Figure 2-Figure Supplement 1*), a canonical member of the nuclear cleavage factor I complex (*Chen and Hyman, 1998*). However, this protein has been shown to shuttle to the cytoplasm and where it may play a regulatory role (*Kessler et al., 1997; Kebaara, 2003; Guisbert et al., 2005*). On average, destabilized mRNAs are longer and contain more optimal codons (*Figure 2-Figure Supplement 6, Khong et al. (2017*)). Together, this analysis suggests that the mechanism of destabilization may act through cis elements in the 5’ UTR and or biased codon usage.

### A genome-wide screen for trans-factors regulating *GAP1* mRNA repression

We sought to identify *trans*-factors mediating accelerated mRNA degradation in response to a nitrogen upshift. We selected *GAP1* as representative of transcript destabilization, as it is abundant in nitrogen-limiting conditions and is rapidly cleared upon addition of glutamine (3.24-fold increase in degradation rate, *Figure 3a, Figure 2-Figure Supplement 12*). Previous approaches to high-throughput genetics of transcriptional activity have used protein expression reporters (*Neklesa and Davis, 2009; Sliva et al., 2016*) or automation of qPCR (*Worley et al., 2015*). However, for our purposes, we required direct measurement of *GAP1* mRNA changes on a rapid timescale. Therefore, we applied single molecule fluorescent *in situ* hybridization (smFISH) to quantify native *GAP1* transcripts in yeast cells in the pooled prototrophic yeast deletion collection (*VanderSluis et al., 2014*). Using fluorescence activated cell sorting (FACS) and Barseq (*Smith et al., 2009; Robinson et al., 2013; Giaever and Nislow, 2014*), we aimed to quantify and model the distribution of *GAP1* mRNA in each mutant (*Kinney et al., 2010; Peterman and Levine, 2016*).

We found that individually labeled probes tiled across *GAP1* mRNA (*Raj et al., 2008*) were insufficiently bright for *GAP1* mRNA quantification using flow cytometry (data not shown), likely due to the small cell size of nitrogen-limited cells and the low transcript numbers in yeast cells compared to mammalian cells (*Klemm et al., 2014*). Therefore, we used branched DNA probes (Quantigene), which serve to amplify the FISH signal (*Hanley et al., 2013*). We developed a fixation and permeabilization protocol (*Figure 4-Figure Supplement 5*) that enabled detection of the distribution of *GAP1* mRNA in steady-state nitrogen-limited conditions and its repression following the upshift (*Figure 3b*). In control experiments, we found that the signal is eliminated in a *GAP1* deletion or by omitting the targeting probe(Fi’gure 3b and *Figure 3-Figure Supplement 1*). To validate sorting, we sorted a sample of cells into quartiles and used microscopy to count fluorescent foci per cell (*Figure 3c*). We found that increased flow cytometry signal is associated with an increase in the number of foci in the cells (*Figure 3d*, R^2^ = 0.607, p < 10^−11^).

**Figure 3.**
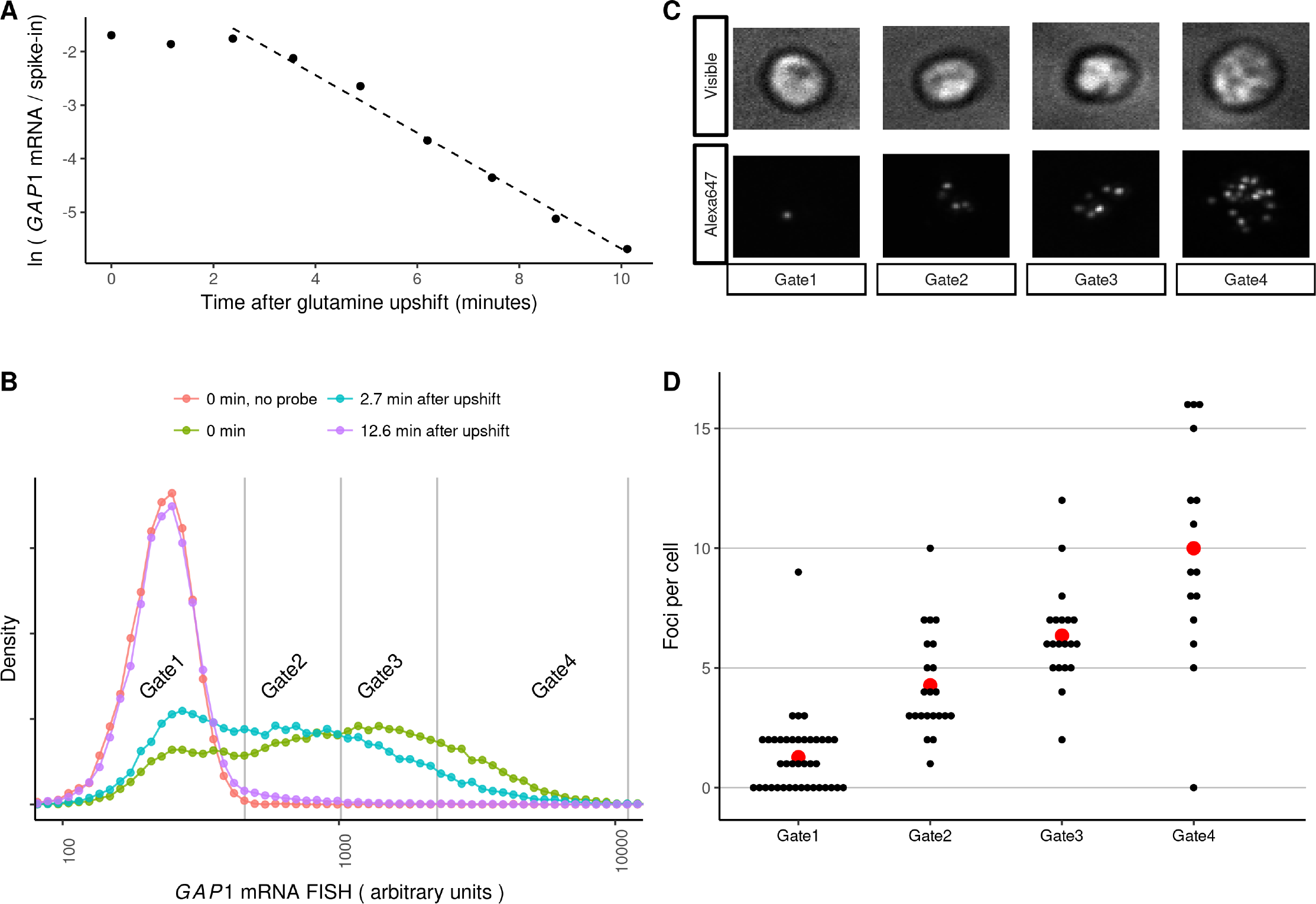
*GAP1* mRNA dynamics measured by flow cytometry. **a)** GAP1 mRNA following upshift measured using RT-qPCR, relative to an external spike-in mRNA standard. The dashed line is fit to points after 2 minutes. **b)** Flow cytometry of wild-type yeast in nitrogen-limited conditions and following an upshift. The vertical grey lines indicate FACS gate boundaries used for cell sorting. **c)** Representative cells from each bin sorted from the experiment in panel b. **d)** Quantification of microscopy data. Each black dot represents a single cell. The mean number of foci per cell in each bin is displayed as a red point. **Figure 3–Figure supplement 1.** GAP1 delete or omission of the targeting probe removes signal of GAP1 FISH.

Previous SortSeq studies of the yeast deletion collection have used outgrowth to generate sufficient material for Barseq (*Sliva et al., 2016*). However, formaldehyde fixation precludes outgrowth. We found that below approximately 10^6^ templates, the Barseq reaction produces primer dimers that outcompete the intended PCR product (*Figure 4-Figure Supplement 5*). Therefore, we re-designed the PCR reaction (*Robinson et al., 2013; Smith et al., 2009*) to be robust for very low sample inputs (*Figure 4-Figure Supplement 5*). Our protocol incorporates a 6-bp unique molecular identifier (UMI) into the first round of extension to identify PCR duplicates, and uses 3’-phosphorylated oligonucleotides and a strand-displacing polymerase (Vent exo-) to block primer dimer formation and off-target amplification. Because strain barcodes are of variable lengths, we developed a bioinformatic pipeline to extract barcodes and UMIs using pairwise alignment to invariant flanking sequences. Based on *in silico* benchmarks, this approach was robust to systematic and simulated random errors that can confound analysis of the yeast deletion barcodes (*Appendix 2, Figure 4-Figure Supplement 5*).

We refer to this experimental approach as BFF (Barseq after FACS after FISH). We used BFF to estimate *GAP1* mRNA abundance for every mutant in the haploid prototrophic deletion collection (*VanderSluis et al., 2014*) in nitrogen-limiting conditions and 10 minutes following the upshift. This approach facilitates identification of mutants with defects in mRNA regulation at both the transcriptional and post-transcriptional level without altering *GAP1* mRNA cis-elements that may affect its regulation. Moreover, this design enables identification of factors that regulate both the steady-state abundance of *GAP1* mRNA and its transcriptional repression following an upshift. We analyzed the deletion pool in biological triplicate (*Figure 4a*). We found that UMIs approached saturation at a slower rate than expected for random sampling, consistent with PCR amplification bias (*Figure 4-Figure Supplement 3*), and therefore we adopted the correction of *Fu et al. (2011*). After filtering, we calculated a pseudo-events metric that approximates the number of each mutant sorted into each bin. Principal components analysis shows that the samples are separated primarily by FACS bin within each condition and biological replicates are clustered indicating that our approach reproducibly captures the variation of *GAP1* mRNA flow cytometry signal across the library (*Figure 4-Figure Supplement 2*).

**Figure 4.**
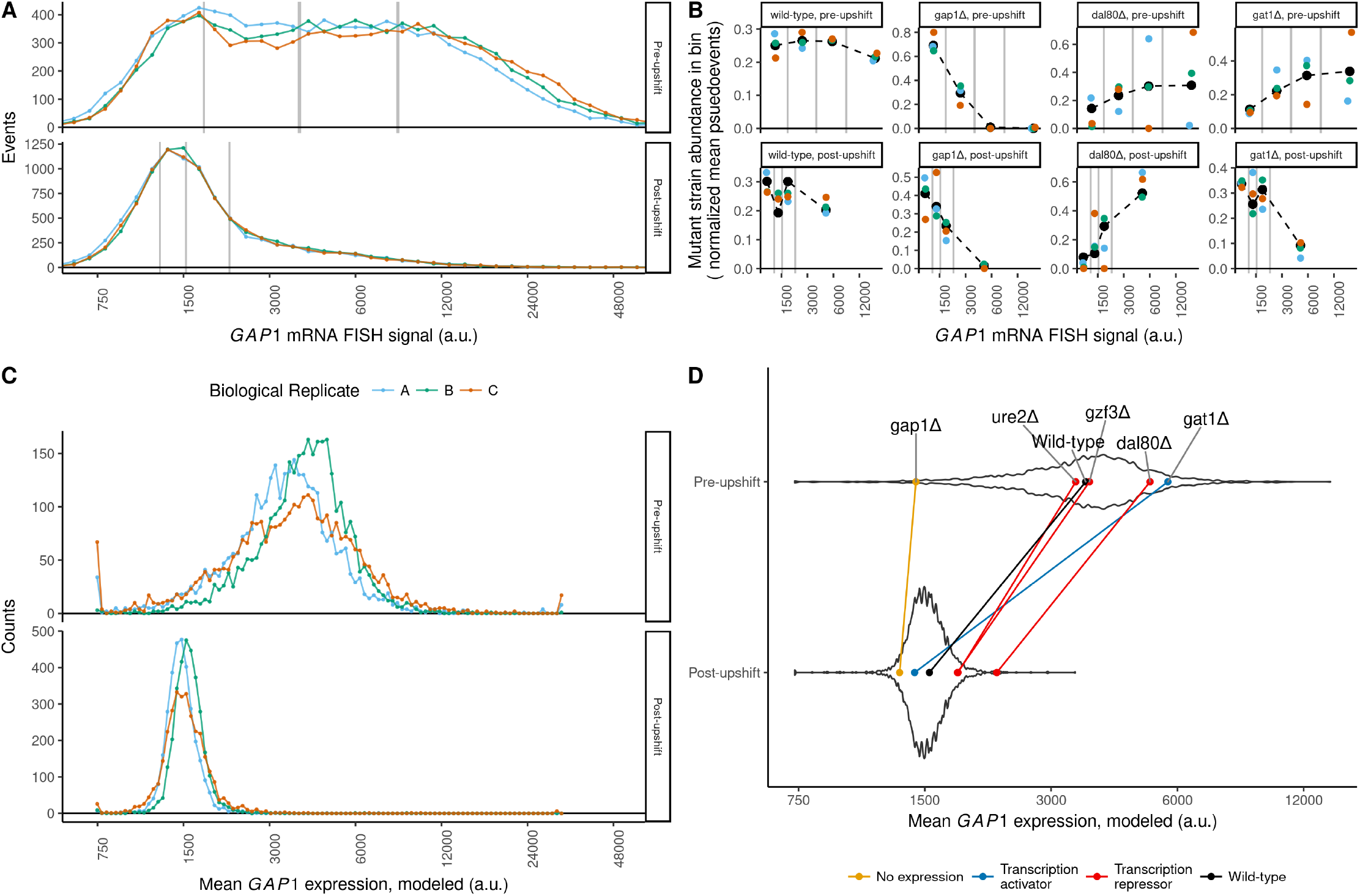
BFF estimates of *GAP1* mRNA abundance. **a)** Flow cytometry analysis of *GAP1* mRNA abundance in the prototrophic deletion collection before and after the upshift. The vertical gray lines denote FACS gates. Biological replicates are indicated by color. **b)** Measurements for individual genes before and after the upshift. Black dashed lines indicate maximum-likelihood fits of a log-normal to pseudo-events for each mutant. Colors and axes as in panel a. **c)** Distribution of mean modeled GAP1 mRNA levels for each mutant. **d)** The mean *GAP1* mRNA expression levels for individual mutants before and after the upshift are shown as points connected by a line, colored according to the type of gene. For reference, the background violin plot shows the distribution of all 3,230 mutants fit. **Figure 4-Figure supplement 1.** Strain barcodes show no length bias, but do show a slight but complex relationship between counts and GC-content. **Figure 4-Figure supplement 2.** Principal components analysis of the abundance estimates for samples. **Figure 4-Figure supplement 3.** Rarefaction curve of total UMI counts against unique UMI counts. **Figure 4-Figure supplement 4.** *tco89*Δ and *xrn1*Δ show defects in *GAP1* mRNA regulation in the BFF assay. **Figure 4-Figure supplement 5.** Supplementary file with experimental rationale, details, and protocol for the BFF experiment. **Figure 4-Figure supplement 6.** Raw counts of strain barcode quantification within each bin in the BFF experiment, and gate settings for the observations. **Figure 4-Figure supplement 7.** BFF data filtered for modeling. **Figure 4-Figure supplement 8.** The parameters of all models fit to the BFF data. **Figure 4-Figure supplement 9.** All 3230 models used for identifying strains with defective *GAP1* dynamics. **Figure 4-Figure supplement 10.** Gene-set enrichment analysis results using *GAP1* estimates.

### Estimating *GAP1* mRNA abundance for individual mutants

We estimated the distribution of *GAP1* mRNA for each mutant by modeling pseudo-events in each quartile as a log-normal distribution using likelihood maximization (*Figure 4b*). From model fits we estimated the mean expression value for each mutant and found that the distribution of means estimated for 3,230 strains (*Figure 4-Figure Supplement 9, Figure 4c*) recapitulates the overall distribution of flow cytometry signal (*Figure 4a*). To validate our approach we first examined strains for which we expected to have a specific phenotype and compared their mean expression level to the distribution of expression for the entire population (*Figure 4d*). We found that the wildtype genotype (*his3Δ,* complemented by the spHis5 in library construction) has an expression level that is centrally located in the distribution both before and following the upshift. The *gap1*Δ genotype is a negative control and we estimate that it is at the extreme low end of the distribution before and following the upshift. *dal80*Δ is a direct transcriptional repressor of NCR transcripts and we found that this is defective in repression of *GAP1* before and after the upshift. Counter-intuitively, deletion of *GAT1,* a transcriptional activator of *GAP1,* appears to have higher steady-state expression of *GAP1* mRNA. However, increased expression of *GAP1* mRNA in a *gat1*Δ background has previously been reported (*Scherens et al., 2006*) and is thought to result from the complex interplay of NCR transcription factors on their own expression levels. Data and models for each mutant strain can be visualized in browser using a Shiny appplication (see http://shiny.bio.nyu.edu/users/dhm267/ or *Appendix 1*).

To identify new cellular processes that regulate *GAP1* mRNA abundance, we used gene-set enrichment analysis (*Figure 4-Figure Supplement 10*). Following the upshift we found mutants that maintain high *GAP1* mRNA expression are enriched for negative regulation of gluconeogenesis (*Figure 5-Figure Supplement 3*) and the Lsm1-7p/Pat1p complex (*Figure 5a*). Mutants in the TORC1 signalling pathway were not enriched; however, we found that a *tco89*Δ mutant has greatly increased *GAP1* mRNA expression before and after the upshift (*Figure 5-Figure Supplement 4*), consistent with the repressive role of TORC1 on the NCR regulon. To compare expression before and after the upshift for each mutant, we regressed the post-upshift mean expression against the pre-upshift mean expression for each genotype (*Figure 5-Figure Supplement 5*). We used the residuals for each strain to identify mutants that clear *GAP1* mRNA with kinetics slower than expected by this model. We found that the Lsm1-7p/Pat1p complex is again strongly enriched for slower than expected *GAP1* mRNA clearance (*Figure 4-Figure Supplement 9*). Specifically the *lsm1 A, lsm6A,* and *pat1*Δ strains are highly elevated in *GAP1* expression before the upshift and strongly impaired in the repression of *GAP1* mRNA after the upshift (*Figure 5a*).

**Figure 5.**
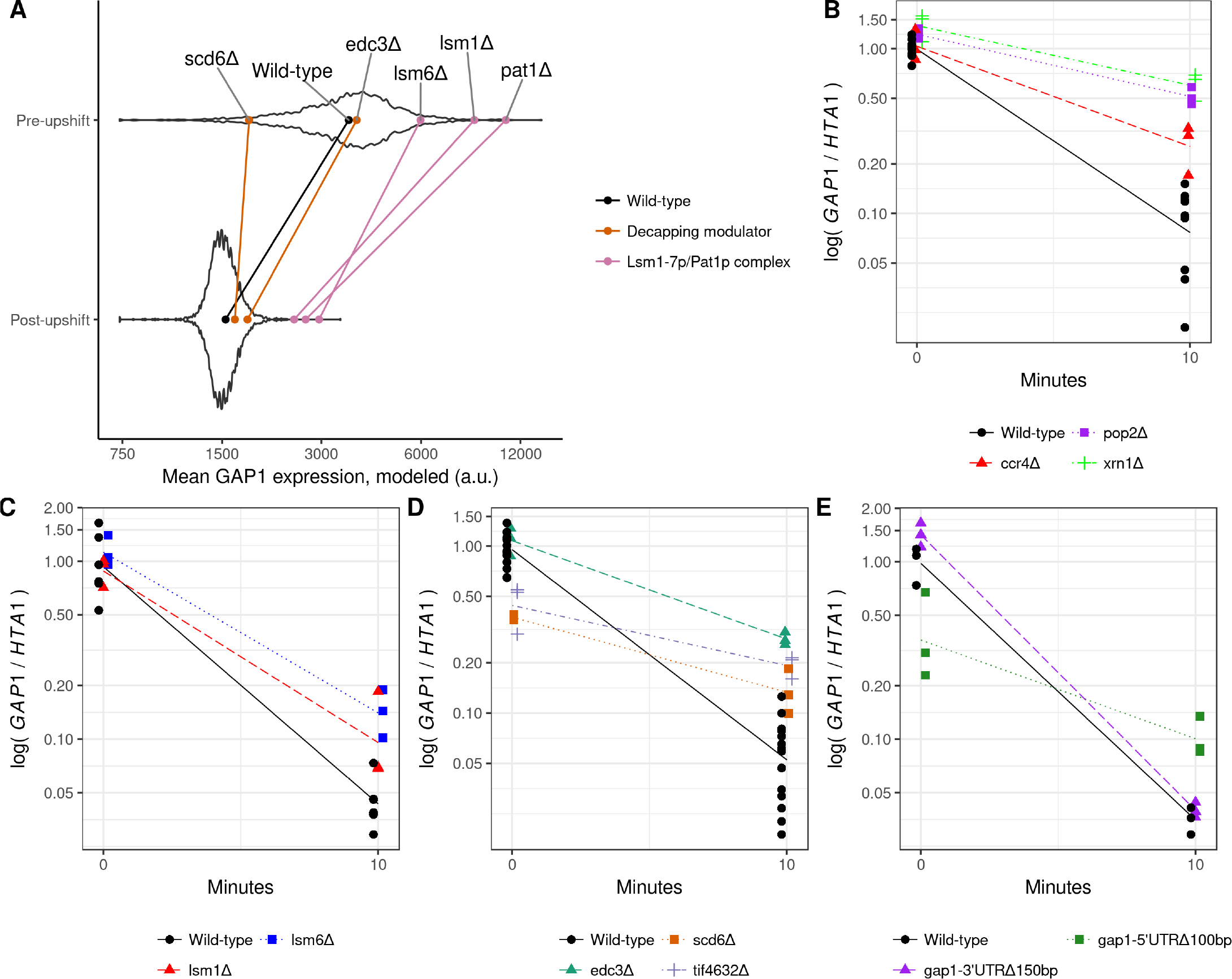
Disrupting the Lsm1-7p/Pat1p complex impairs clearance of *GAP1* mRNA. **a)** In the background is the distribution of fit GAP1 mRNA mean expression levels for all mutants in the pool. Indicated by colored points and lines are the means for individual knockout strains, as labeled. **b-e**), *GAP1* mRNA relative to *HTA1* mRNA before and 10 minutes after a glutamine upshift, in biological triplicates. Lines are a log-linear regression fit. Points are dodged horizontally for clarity, but timepoints for modeling and for drawn lines are 0 and 10 minutes exactly. Wild-type is FY4. **b)** xrn1Δ, ccr4Δ, pop2Δ are all slowed in clearance (p-values < 0.004). c) lsm1Δ and lsm6Δ are slowed in clearance (p-values < 0.0132 and 0.0299, respectively). **d)** edc3Δ is slowed in clearance (p-value < 10^−4^). scd6Δ and tif4632Δ are slowed in clearance (p-values < 10^−5^) and have lower levels of expression before the upshift (p-values < 0.003). **e)** A deletion of 150bp 3’ of GAP1 stop codon has no significant effect, but a deletion of 100bp 5’ of the start codon has slower clearance (p-value < 10^−4^) and lower level of expression before the upshift (p-value < 0.0015). **Figure 5–Figure supplement 1.** An independently generated GAP1 5’ deletion. **Figure 5–Figure supplement 2.** Processing-body dynamics are not associated with the nitrogen upshift, by Dcp2p-GFP microscopy. **Figure 5–Figure supplement 3.** Knock-out mutants of negative regulators of gluconeogenesis are associated with higher GAP1 expresion after the upshift. **Figure 5–Figure supplement 4.** Knock-out mutants of genes involved in sulfate assimilation are associated with higher estimated GAP1 mean after the upshift. **Figure 5–Figure supplement 5.** The relationship between the estimated mean before the shift and after the upshift.

As these factors are associated with processing-body dynamics, we tested if microscopically-observable processing-bodies form or disassociate during the upshift, using microscopy of Dcp2-GFP. We did not observe qualitative changes in Dcp2-GFP distribution (*Figure 5-Figure Supplement 2*), and thus the upshift does not result in a microscopically visible changes in processing-body foci as seen in other stresses. This is consistent with previous investigations of amino-acid limitation stress (*Hoyle et al., 2007*) and suggests that the defects in *GAP1* mRNA clearance likely result from their roles in decapping or associated processes.

To confirm the role of the Lsm1-7p/Pat1p complex in clearing *GAP1* mRNA during the nitrogen upshift we measured *GAP1* mRNA repression using qPCR normalized to *HTA1,* which is not subject to destabilization upon the upshift (*Figure 2a*). We also tested mutants that were not detected using BFF, or were only detected in the highest *GAP1* bin and therefore not suitable for modeling (e.g. *xrn1A Figure 5-Figure Supplement 4*). Using this assay we found that the main 5’-3’ exonuclease *xrn1*Δ and mRNA deadenylase complex (ccr4A and *pop2*Δ) are impaired in *GAP1* repression (*Figure 5b*). We found that qPCR confirms results from BFF. We confirmed that the accelerated degradation of *GAP1* mRNA is impaired in *lsm1*Δ and *lsm6A (Figure 5c*). We also tested *scd6*Δ and *edc3Δ,* two modifiers of the decapping or processing-body assembly functions associated with this complex, and found two distinct phenotypes (*Figure 5d*). *edc3*A has similar expression as wild-type before the upshift, but is cleared much more slowly. *scd6*A has a greatly reduced *GAP1* expression before the upshift but is impaired in *GAP1* clearance. *tif4632Δ,* a deletion of the eIF4G2 known to interact with Scd6p (*Rajyaguru et al., 2012*), has a similar phenotype.

Identification of an initiation factor subunit with defects in *GAP1* mRNA clearance suggests that translation control may impact stability changes. Therefore we deleted the 5’ UTR and 3’ UTR of *GAP1*. Whereas the 3’ UTR deletion does not have an effect the 5’ UTR deletion exhibit the phenotype of reduced *GAP1* mRNA before the upshift and a reduced rate of transcript clearance following the upshift (*Figure 5e*). We observed a similar phenotype with a different deletion of 152bp upstream of the *GAP1* start codon (*Figure 5-Figure Supplement 1*). This indicates that cis-elements responsible for the rapid clearance of *GAP1* are unlikely to be located in the 3’ UTR, and instead may be exerting an effect at the 5’ end of the mRNA.

## Discussion

Regulated changes in mRNA stability allows cells to rapidly reprogram gene expression, clearing extant transcripts that are no longer required and potentially reallocating translational capacity. Pioneering work in budding yeast has shown that mRNA stability changes facilitate gene expression remodeling in response to changes in nutrient availability including changes in carbon sources (*Scheffler et al., 1998*) and iron starvation (*Puig et al., 2005*). Here, we characterized genome-wide changes in mRNA stability in response to changes in nitrogen availability and identified factors that mediate the rapid repression of the destabilized mRNA, *GAP1.* Our study extends our previous work characterizing the dynamics of transcriptome changes using chemostat cultures (*Airoldi et al., 2016*) and shows that accelerated mRNA degradation targets a specific subset of the transcriptome in response to changes in nitrogen availability. We developed a novel approach to identify regulators of mRNA abundance using pooled mutant screens and find that modulators of decapping activity, and core degradation factors, are required for accelerated degradation of *GAP1* mRNA.

Measuring the stability of the transcriptome requires the ability to separate pre-existing and newly synthesized transcripts. We modified existing methods to measure post-transcriptional regulation of the yeast transcriptome in a nitrogen upshift using 4-thiouracil labeling (*Miller et al., 2011; Neymotin et al., 2014; Munchel et al., 2011*). These modifications entailed improved normalization and quantification of extant transcripts and explicit modeling of labelling dynamics to account for some of the inherent limitations of metabolic labeling approaches (*Pérez-Ortín et al., 2013*). Continued development of fractionation biochemistry (*Duffy et al., 2015*) and incorporation of explicit per-transcript efficiency terms will improve these methods further (*Chan et al., 2017*).

Our experiments show that the process of physiological and gene expression remodeling occur on very diferent timescales in response to a nitrogen upshift. Cellular physiology is remodeled over the course of two hours to achieve a new growth rate. By contrast, transcriptome remodeling occurs rapidly and through states that are distinct from increases in steady-state growth rates. Previous studies have shown that transcriptional activation of the NCR regulon is rapidly repressed upon a nitrogen upshift (*Airoldi et al., 2016*). Our results indicate that accelerated degradation of many NCR transcripts (*Godard et al., 2007*) contributes to this repression. A three-fold increase in the degradation rate of *GAP1* mRNA provides an additional layer of repressive control. Importantly, our results show that accelerated degradation is not limited to NCR transcripts but also targets transcripts enriched in carbon metabolism pathways, particularly pyruvate metabolism. Conversely, we also detect an apparent reduction in the degradation rate for some transcripts including *MAE1. MAE1* encodes an enzyme responsible for the conversion of malate to pyruvate, and combined with the accelerated degradation of *PYK2* mRNA may reflect an adaptive shunt of carbon skeletons from glutamine to alanine via the TCA cycle (*Boles et al., 1998*). Recently, *Tesnière et al. (2017)* described destabilization of carbon metabolism mRNAs after repletion of nitrogen following 16 hours of starvation. We do not detect destabilization of *PGK1* mRNA and note that the basal half-life of 6.2 minutes estimated in our study is similar to the accelerated rate reported by *Tesnière et al. (2017)*.

Destabilized transcripts are enriched for a binding motif of Hrp1p in the 5’ UTR. This essential component of mRNA cleavage for poly-adenylation in the nucleus has been shown to shuttle to the cytoplasm and bind to amino-acid metabolism mRNAs (*Guisbert et al., 2005*) and been shown to interact genetically to mediate nonsense-mediated decay (NMD) of a *PGK1* mRNA harboring a premature stop-codon (*González et al., 2000*) or a cis-element spanning the 5’ UTR and first 92 coding bp of *PPR1* mRNA (*Kebaara, 2003*). A potential role for these Hrp1p sites warrants further investigation.

BFF identified mutants in the Lsm1-7p/Pat1p complex as having elevated *GAP1* mRNA levels both before and after the upshift. This is expected given their central role in mRNA metabolism, and experiments using *GAP1* normalized to *HTA1* demonstrate that the effect before the upshift is likely a global effect (*Figure 5c*). However, these mutants still have a significant defect in clearance of *GAP1,* and the assay demonstrates that associated decapping factors *EDC* and *SCD6* have specific effects (*Figure 5d*). Given that the *GAP1* mRNA is destabilized during this transition we suspect that these mRNA degradation factors are directly involved. While we found that the *edc3*Δ mutant has defects in clearance of *GAP1,* we also found that scd6A, *tif4632Δ,* and deletion of the 5’ UTR of *GAP1* impairs clearance (*Figure 5e*). This deletion does not include the TATA box (ending at -179) or GATAA sites (nearest at -237) responsible for NCR GATA-factor regulation of *GAP1 (Stanbrough and Magasanik, 1996*). This suggests that interactions of these factors with cis-elements in the 5’ UTR might be responsible for stabilizing *GAP1* mRNA during limitation, although the truncation of the 5’ sequence may be enough to inhibit translation initiation by virtue of the shorter length (*Arribere and Gilbert, 2013*). Elements in the 5’ UTR have also been demonstrated to modulate *GAL1* mRNA stability (*Baumgartner et al., 2011*) and destabilize *SDH2* mRNA upon glucose addition, perhaps due to the competition between translation initiation and decapping mechanisms (*de la Cruz et al., 2002*). Interestingly, both *GAP1* and *SDH2* share the feature of a second start codon downstream of the canonical start (*Neymotin et al., 2016*) and we have previously found that mutation of the start codon of *GAP1* results in lower steady-state mRNA abundances (*Neymotin et al., 2016*). This suggests a mechanism of degradation through dynamic changes in translation initiation that triggers decapping of *GAP1* and other mRNA. Future work interrogating this possible interaction of translational status and mRNA stability during dynamic conditions could also expand our understanding of the relationship between these two processes.

To our knowledge, this is the first time mRNA abundance has been directly estimated using a SortSeq approach, although using mRNA FISH and FACS to enrich subpopulations of cells has been previously reported (*Klemm et al., 2014; Hanley et al., 2013; Sliva et al., 2016*). This approach could be used with other barcoding mutagenesis technologies, like transposon-sequencing or Cas9 mediated perturbations, to systematically test the genetic basis of transcript dynamics. The use of branched-DNA mRNA FISH, or other methods (*Rouhanifard et al., 2017*), allows for mRNA abundance estimation without requiring genetic manipulation which makes it suitable for a variety of applications such as extreme QTL mapping. Furthermore, our methods for library construction should permit accurate quantification of pooled barcode libraries with small inputs, expanding the possibilities for flow cytometry markers to fixed-cell assays.

Why is *GAP1* subject to multiple layers of gene product repression upon a nitrogen upshift, at the level of transcript synthesis, degradation, protein maturation, and post-translational inactivation? Given the strong fitness cost of inappropriate activity (*Risinger et al., 2006*), this overlap could ensure mechanistic redundancy for robust repression in the face of phenotypic or genotypic variation. Alternatively, it could reflect a systematic need to free ribonucleotides or translational capacity, or result from some as yet uncharacterized process. Future work aimed at determining the adaptive basis of accelerated mRNA degradation will serve to illuminate the functional role of post-transcriptional gene expression regulation.

## Methods and Materials

### Availability of data and analysis scripts

Computer scripts used for all analyses are available as a git repository on GitHub (https://github.com/darachm/millerBrandtGresham2018) and data is available as zip archives on the Open Science Framework (https://osf.io/hn357/). Instructions for obtaining, unpacking, and using these are in *Appendix 2.*

We did not make this 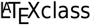, it is a modified version of one provided by a journal.

### Media and upshifts of media

Nitrogen-limited media (abbreviated as “Nlim”) is a minimal media supplemented with various salts, metals, minerals, vitamins, and 2% glucose, as previously described (*Airoldi et al., 2016; Brauer et al., 2008*). For proline limitation, Nlim base media was made with 800*μ*M L-proline as the sole nitrogen source (NLim-Pro). YPD media was made using standard recipes (*Amberg et al., 2005*). All growth was at 30°C, in an air-incubated 200rpm shaker using baffled flasks with foil caps, or roller drums for overnight cultures in test tubes. For glutamine upshift experiments, 400*μ*M L-glutamine was added from a 100mM stock solution dissolved in MilliQ double-deionized water and filter sterilized. All upshift experiments were performed at a cell density between 1 and 5 million cells per mL, in media where saturation is approximately 30 million cells per mL. For all experiments, a colony was picked from a YPD plate and grown in a 5mL NLimPro pre-culture overnight at 30^°^C, then innoculated into the experimental culture from mid-exponential phase.

### Strains

See *Table 2* for details. The wild-type strain used is FY4, a S288C derivative. The pooled deletion collection is as published in *VanderSluis et al. (2014*). For all experiments with single strains, colonies were struck from a -80°C frozen stock onto YPD (or YPD+G418 for deletion strains) to isolate single colonies. For pooled experiments we inoculated directly into NLim media from aliquots of frozen glycerol stocks.

**Table 2.** Yeast strains used in this study

| Strain ID | Short description | Details |
| --- | --- | --- |
| DGY1 | FY4 | Isogenic to S288C, prototrophic, MATa |
| - | Deletion collection pool | Haploid (MATa) prototrophic deletion collection as described in the publication of **VanderSluis et al. (2014)** |
| DGY410 | xrn1Δ::KanMX | ygl173cΔ::KanMX from the prototrophic deletion collection |
| DGY564 | ccr4Δ::KanMX | yal021cΔ::KanMX from the prototrophic deletion collection |
| DGY565 | pop2Δ::KanMX | ynr052cΔ::KanMX from the prototrophic deletion collection |
| DGY547 | lsm1Δ::KanMX | yjl124cΔ::KanMX from the prototrophic deletion collection |
| DGY571 | lsm6Δ::KanMX | ydr378cΔ::KanMX from the prototrophic deletion collection |
| DGY545 | pat1Δ::KanMX | ycr077cΔ::KanMX from the prototrophic deletion collection |
| DGY554 | edc3Δ::KanMX | yel015wΔ::KanMX from the prototrophic deletion collection |
| DGY552 | scd6Δ::KanMX | ypr129wΔ::KanMX from the prototrophic deletion collection |
| DGY611 | tif4632Δ::KanMX | ygl049cΔ::KanMX from the prototrophic deletion collection |
| DGY539 | GAP1 5' UTR delete | confirmed by Sanger sequencing to have 152bp deleted 5' of the start codon |
| DGY576 | GAP1 5' UTR delete | confirmed by Sanger sequencing to have 100bp deleted 5' of the start codon |
| DGY577 | GAP1 3' UTR delete | confirmed by Sanger sequencing to have 150bp deleted 3' of the stop codon |
| DGY525 | FY3 + pRP1315 | FY3, a ura- auxotroph (ura3-52), transformed with pRP1315 (URA3 marker, expressing a Dcp2-GFP fusion) |

Strains with deletions 5’ of the start codon and 3’ of the stop codon were generated by the “delitto-perfeto” method (*Storici and Resnick, 2006*), by inserting the pCORE-Kp53 casette at either the 5’ or 3’ end of the coding sequence, then transforming with a short oligo product spanning the deletion junction and counter-selecting against the casette with Gal induction of p53 from within the cassette. These strains were generated and confirmed by Sanger sequencing, and traces are available in directory data/qPCRfoiiowup/ within the data zip archive (*Appendix 2*).

### Measurement of growth during upshift

A single colony of FY4 was inoculated in 5mL NLimPro media and grown to exponential phase, then back diluted in NLimPro media in a baffled flask. Samples were collected into an eppendorf, sonicated, diluted in isoton solution, and analyzed with a Coulter Counter Z2 (Beckman Coulter).

### Re-analysis of microarray data

Gene expression data (*Airoldi et al., 2016*) were analyzed using pcaMethods to perform a SVD PCA on scaled data.

### qPCR

Each strain was grown from single colonies. Samples were collected before, during the first ten minutes of the nitrogen upshift (*Figure 3*), or at ten minutes after the upshift (*Figure 3*). For the experiments described in *Figure 5,* all work was done in biological replicates. Each 10mL sample was collected by vacuum onto a 25mm nylon filter and frozen in an eppendorf in liquid nitrogen. RNA was extracted by adding 400*μ*L of TES buffer (10mM Tris (7.5pH), 10mM EDTA, 0.5% SDS) and 400*μ*L of acid-phenol, vortexing vigorously and incubating at 65°C for an hour with vortexing every 20 minutes. For *Figure 3* only, at the beginning of this extraction incubation we added 10*μ*L of a 0.1ng/*μ*L in-vitro synthesized spike-in mRNA BAC1200 (as generated for the label-chase RNAseq (*Figure 2-Figure Supplement 7*), but without 4-thiouridine). All samples were separated by centrifugation and extracted again with chloroform on a 2mL phase-lock gel tube (5Prime #2302830). After ethanol precipitation of the aqueous layer, RNA was treated DNAse RQ1 (Promega M610A) according to manufacturer instructions, then the reaction heat-killed at 65°C for 10 minutes after adding a mix of 1:1 0.5M EDTA and RQ1 stop-solution. The resulting RNA was cleaned with a phenol-chloroform extraction and ethanol precipitated. All samples were hybridized with RT primers by incubating the mixture at 80° for 5 minutes then on ice for 5 minutes. For *Figure 3* 2*μ*g RNA was primed with 2.08ng/*μ*L random hexamers (Invitrogen 51709) and 2.5mM total dNTPs (Promega U1511), while for *Figure 5* 1*μ*g RNA was primed with 5.6mM Oligo(dT)18 primers (Fermentas FERSO132) and 0.56mM total dNTPs (Promega U1511). These mixtures were combined with 1/10th 10x M-MulvRT buffer (NEB M0253L), 1/20th volume RNAse-OUT (Invitrogen 51535), and 1/20th volume M-MulvRT (NEB M0253L). A negative control with no reverse-transcriptase enzyme was also prepared and analyzed in the qPCR reaction. The reaction proceeded for 1 hour at 42^°^C, then was heat-killed at 90^°^C before diluting 1/8 with hyclone water (GE SH30538). This dilution was used as direct template in 10*μ*L reactions with SybrGreen I Roche qPCR master-mix (Roche 04 707 516 001) for measurement on a Roche Lightcycler 480. For *Figure 3,* we used primers DGO230,DGO232 to quantify *GAP1* and DGO605,DGO606 to quantify the synthetic spike-in BAC1200. For *Figure 5,* we used primers DGO229, DGO231 to quantify *GAP1* and DGO233, DGO236 to quantify *HTA1.* See *Table 3* for sequence. These were run on a Roche480 Lightcycler, with a max-second derivative estimate of the cycles-threshold (the *c* value output by analysis) used for analysis by scripts included in the git repo (*Appendix 2*). Linear regression of the log-transformed values was used to quantify the dynamics and assess significance of changes in expression levels or rates of change.

**Table 3.**
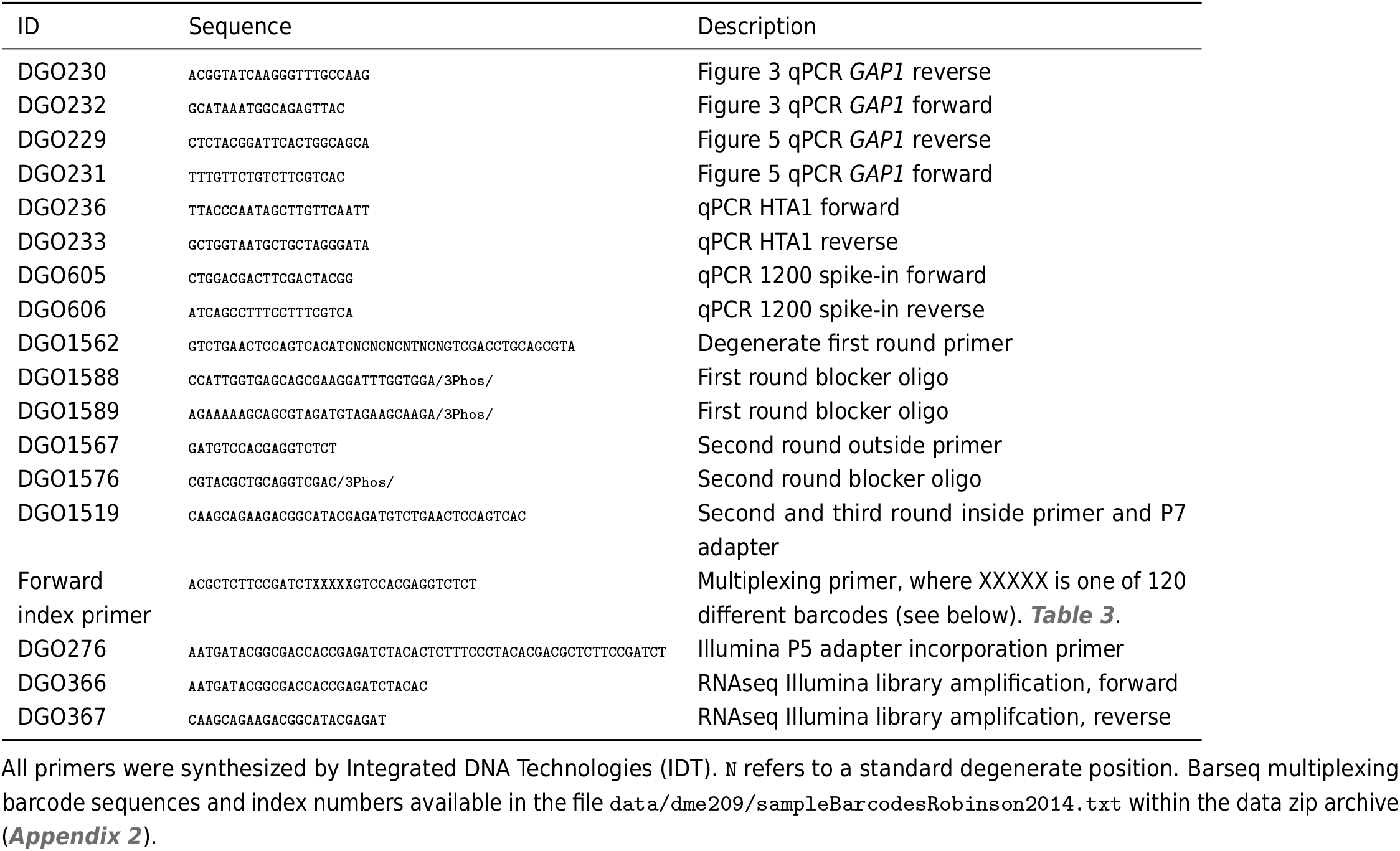
Primers used in this study

### Microscopy of Dcp2-GFP

To look for processing-body dynamics in response to a nitrogen upshift, we used strain DGY525, which is FY3 containing plasmid pRP1315 (gift from Roy Parker). Samples were collected before and following a nitrogen upshift, from exponential growth in YPD, or 10 minutes after resuspending YPD-grown cells in DI water. All samples were collected by centrifugation at 10,000g for 30 seconds, aspirating most supernatant, then centrifugation for 20 seconds and aspirating all media. Each pellet was immediately resuspended in 4% PFA (diluted from EMS 16% PFA ampule RT15710) with 1x PBS ( NaCl 8g/L, KCl 0.2g/L, Na_2_HPO_4_ 1.42g/L, KH_2_PO_4_ 0.24g/L) for 4, 10, 12, 19, or 25 minutes on bench, then spun at 10,000g for 1 minute, aspirated, then washed once and resuspended with 1x PBS. Samples were kept on ice, then put onto a coverslip for imaging on a DeltaVision scope. Raw images available in the microscopy zip archive (*Appendix2*).

### 4tU label-chase and RNA sequencing

The methods and analysis are detailed in *Figure 2-Figure Supplement 7,* including design rationale, protocols, and manufacturer information, and all data and code are available according to instructions in *Appendix 2.*

FY4 was grown in nitrogen-limitation conditions overnight with a 50*μ*M:50*μ*M mixture of 4-thiouracil:uracil. This culture was split, then 4mM uracil was added to chase the 4-thiouracil label (a 41-fold excess of uracil). 30mL samples of the culture were taken by filtration onto 25mm nylon filters and flash-frozen in eppendorfs. After letting the chase proceed for 12.5 minutes, we added glutamine from a 100mM stock (dissolved in water) to a final concentration of 400*μ*M to one flask, or an equal amounts of water to the control flask. Samples were extracted using a hot acid-phenol method, with equal volume of synthetic spike-ins added to each RNA extraction reaction. 4tU-containing spike-ins (polyadenylated coding sequences from *B. subtilus* and *C. elegans*) were synthesized *in-vitro* as previously described (*Neymotin et al., 2014*). RNA was reacted with MTSEA-biotin to conjugate biotin to the 4-thiouracil-containing RNA, then purified using streptavidin beads. Fractionated RNA was depleted of rRNA using a RiboZero kit. RNA samples were converted into Illumina sequencing libraries using a strand-specific (UNG) protocol, ligating adapters that contain UMI’s (*Hong and Gresham, 2017*). These libraries were pooled and sequenced by the NYU Genomics Core sequencing facility on an Illumina NextSeq. Following base-calling and sample demultiplexing by NYU GenCore, the sequencing reads were trimmed using cutadapt (*Martin, 2011*) aligned using tophat2 (*Kim et al., 2013*) to a reference genome that included the yeast reference genome (assembly R64) and spike-ins, filtered for mapping-quality and length using `samtoois` (*Li et al., 2009*), deduplicated with `umi_toois` (*Smith et al., 2017*) and feature counting was performed using `htseq-count` (*Anders et al., 2015*). Feature counts for yeast mRNAs were normalized to synthetic spike-ins, using the fitted values from a log-linear model of spike-in abundance increase (see Results, and *Figure 2-Figure Supplement 7*). The rate of mRNA degradation and changes in this rate was quantified to assume an exponential model *Figure 2-Figure Supplement 7,* fit as a linear model to log transformed data. Significant changes in mRNA degradation rates were quantified from the coefficient method of the linear model using a cut-off of a FDR (*Storey et al., 2015*) less than 0.01 and a doubling in degradation rate (based on modeling detailed in *Figure 2-Figure Supplement 7*).

### Label-chase RNA sequencing *cis* element analysis

To detect if *de novo* or known *cis* elements were associated with destabilization upon a nitrogen upshift, we used DECOD (*Huggins et al., 2011*), FIRE (*Elemento et al., 2007*), TEISER (*Goodarzi et al., 2012*), and the #ATS pipeline (*Li et al., 2010*). We also scanned for association with RBP binding sites from the CISBP-RNA database (*Ray et al., 2013*) using AME from the MEME suite (*McLeay and Bailey, 2010*). Final plots in the supplement were made using motif scans with GRanges (*Lawrence et al., 2013*). Analysis was done using coding sequence and four different definitions of untranslated regions (200bp upstream of the start codon or downstream of the stop codon, the largest detected isoform in TIF-seq data (*Pelechano et al., 2014*), or the most distal detected gPAR-CliP sites in exponential-phase or nitrogen-limited growth (*Freeberg et al., 2013*)).

### Barseq after FACS after mRNA FISH (BFF)

The methods and analysis are detailed in *Figure 2-Figure Supplement 7,* including design rationale, protocols, and manufacturer information.

An aliquot of the prototrophic deletion collection (*VanderSluis et al., 2014*) was thawed and diluted, with approximately 78 million cells added to 500mL of NLimPro media in a 1L baffled flask. This was shaken at 30°C overnight, then split into three flasks (A, B, and C). After three hours (at mid-exponential) we collected samples of 30mL culture filtered onto a 25mm filter and flash-frozen in an eppendorf in liquid nitrogen. We sampled in steady-state growth (pre-upshift) and 10.5 minutes after adding 400*μ*M glutamine (post-upshift). Samples of the pool were fixed with formaldehyde (4% PFA diluted in PBS from 10mL aliquot, buffered, 2 hours room-temperature) and digested with lyticase (in BufferB with VRC 37° 1 hour), (*McIsaac et al., 2013*), and permeabilized with ethanol at 4° overnight. Samples were processed with a Affymetrix Quantigene Flow RNA kit (product code 15710) designed to target for *GAP1* mRNA and labelled with Alexa 647. This hybridization was done using a modified version of the manufacturer’s protocol (Appendix *Figure 4-Figure Supplement 5,* including a DAPI staining step. Samples were sonicated, then run through a BD FACSAria II. Cells were gated for singlets and DAPI content (estimated 1N or more), then sorted based on emission area from a 660/20nm filter with a 633nm laser activation into four gates within each timepoint, across replicates. These were sorted using PBS sheath fluid at room-temperature, into poly-propylene FACS tubes, then stored at -20°C. For each gate, cells were collected via centrifugation and genomic DNA extracted by NaCl reverse-crosslinking at 65°, inspired by *Klemm et al. (2014*), with subsequent proteinase K and RNase A digestions. Genomic DNA was split into three reactions to amplify in a modified barseq protocol (*Figure 4-Figure Supplement 5*). See the supplementary write-up *Figure 4-Figure Supplement 5* for detailed protocols, rationale, and a discussion of dimers. Barseq libraries were submitted to the NYU Genomics Core for sequencing on a 1x75bp run on a Illumina NextSeq.

### Analysis of BFF sequencing results

We devised a pipeline to quantify barcodes using the UMI sequence incorporated in the first round of amplicon priming, and benchmarked on *in silico* simulated datasets *Figure 4-Figure Supplement 5.* Briefly, raw FASTQ files are processed with SLAPCHOP (https://github.com/darachm/slapchop">) which uses pair-wise alignment (*Cock et al., 2009*) to filter, extract UMIs from variable positions, and extract barcodes into different fields. We demultiplex using a perl script, and align paritioned strain barcodes to a reference barcode index *Smith et al. (2009*) using `bwa mem` *Li (2013*). Barcodes are counted, then we used the UMI’s with the label-collision correction of *Fu et al. (2011*) to quantify the proportion of each mutant in the sample. These relative counts are used the FACS data (the sorted events per bin) to estimate the distribution of each mutant across the four gates in each timepoint. We filtered for strains detected in at least three bins, and fit a log-normal distribution using `mle` in R *Team (2000*). The mean of this distribution was used as the expression value of *GAP1* in plots and GSEA analysis using `clusterProfiler` (*Yu et al., 2012*).

## Acknowledgments

We would like to acknowledge the funding source of NIH grant 5R01GM107466. We would also like to thank Andreas Hochwagen and Viji Subramanian for microscope access, Ken Birnbaum for helpful conversations and equipment usage, Michi Pedraza, Andres Mansisidor, and Matt Paul for helpful reading and comments, Evelina Tutucci for demonstrating FISH, staff at eBioscience/Affymetrix/ThermoFisher for support with the Quantigene/FlowRNA probe set, the Cold Spring Harbor Yeast Course, the NYU Genomics Core facility for sequencing, flow cytometry, and support, and past and present members of the Gresham and Vogel labs for discussions and support.

## Visualizing data with a Shiny application

A Shiny application is available to explore the two main datasets in this paper, at http://shiny.bio.nyu.edu/users/dhm267/. It is also included as a separate zipped archive for local installation and long-term archiving. To use the Shiny applications from the zipped archive:

1. Download independent_shiny_archive.zip.
2. Unzip this archive.
3. Open R.
4. Install the ’shiny’ and ‘tidyverse’ packages by entering the command `install.packages(c(“shiny”,”tidyverse”))`
5. Enter the command: `shiny::runApp(“shiny”,port=5000)` where “shiny” is the path to the unziped folder and “5000” is an arbitrarily selected port number.
6. Point your web browser at the URL 127.0.0.1:5000 and follow the instructions. The application has two tabs, one for the label-chase RNAseq and one for the BFF experiment.

## Organization and availability of code and data

The computer analysis code is available as a git repository on Github:

https://github.com/darachm/millerBrandtGresham2018

The data are available as a set of ‘zip’ format archives on OSF:

https://osf.io/7ybsh/

To reproduce the entire analysis, or to access a particular analysis, clone the git repo. For example, on a Linux/Unix/MacOSX system install ‘git’ and run:

git clone https://github.com/darachm/millerBrandtGresham2018.git

and change into that directory. Then, download the ‘zip’ data archives from the above OSF link, and put them inside this git repo folder (here, ‘millerBrandtGresham2018’). At minimum, you should have the ‘data.zip’ archive in that directory, although records of all R analyses are in ‘html_reports.zip’ and intermediate files are in ‘tmp.zip’.

Consult the ‘README.md’ file in the repository for more instructions and options, including to unzip intermediate files and HTML reports generated for every R script which detail the results.

**Figure 1–Figure supplement 1.**
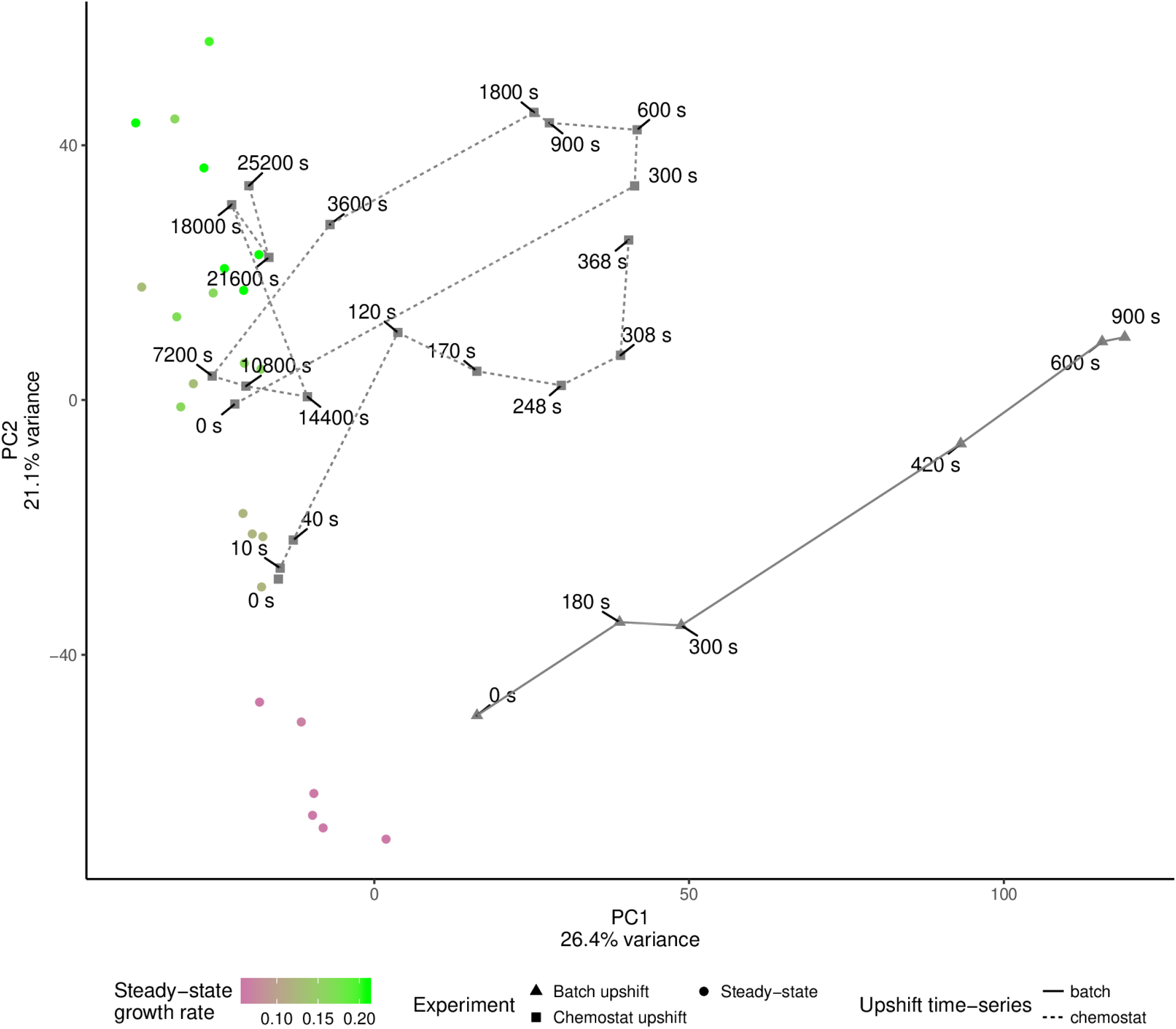
The coarse long-term transcriptome dynamics of a glutamine upshift. Principal components analysis (SVD) of microarray data from Airoldi et al. (2016). Colored points are from steady-state chemostats grown in limitation for various nitrogen sources, at different growth rates. Time-series experiments are show in grey points, connected by lines, and line-type is the type of upshift (in batch or in chemostat).

Figure1_Table_GSEofGOtermsAgainstPCcorrelation.csv

Figure 1-Figure supplement 2. Gene set enrichment analysis of loadings on principal components one and two.

**Figure 2–Figure supplement 1.**
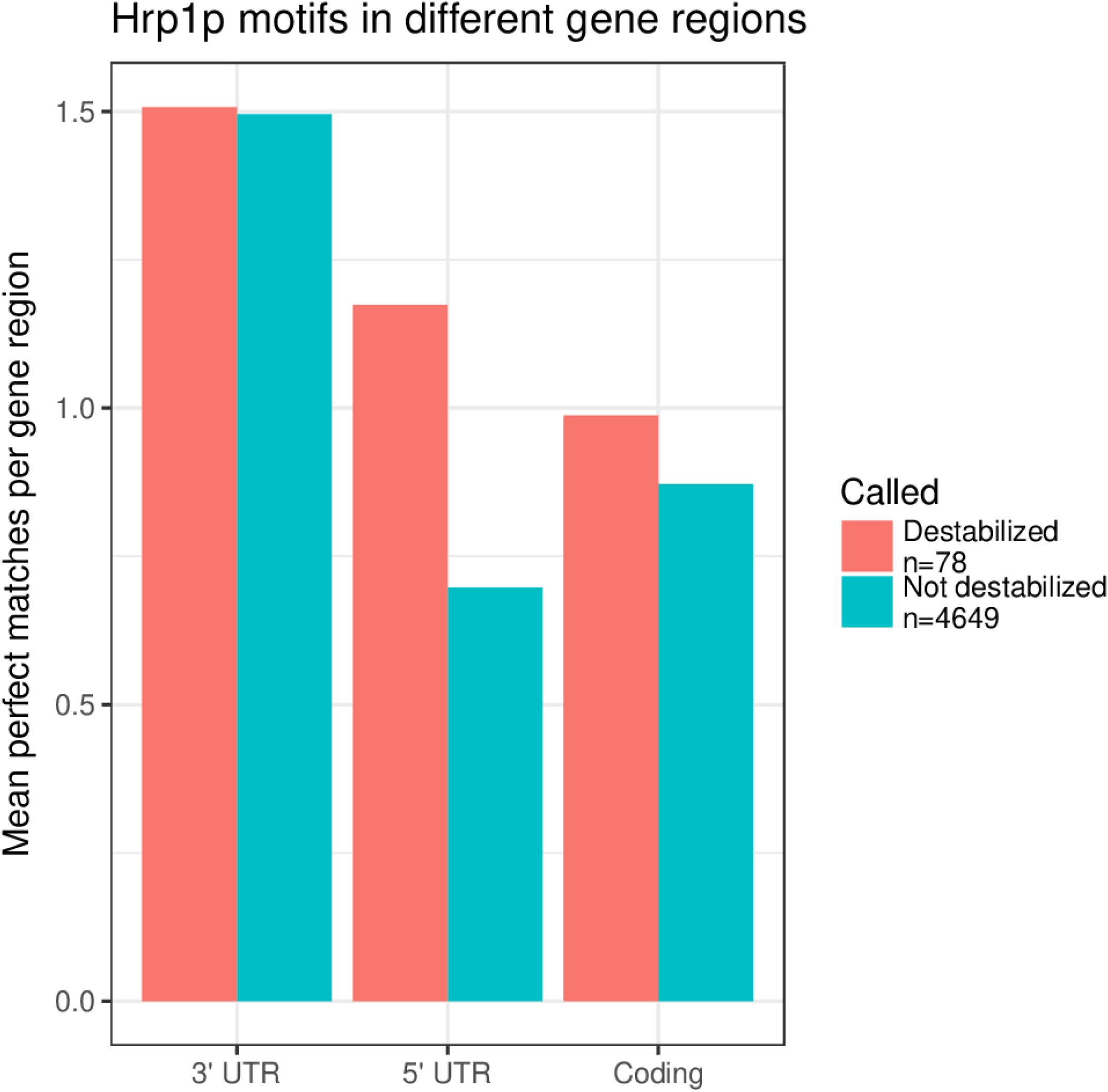
Sequences of destabilized and unaltered mRNAs were analyzed for RBP binding motif enrichment using the AME program in MEME, then significant hits were confirmed by using a logistic model predicting destabilization based on motif content per sequence length. Hrp1p is significantly ( p<0.0001) enriched in the 5’ UTRs of destabilized transcripts. For this plot, motif matches were counted using the GRanges package for the 5’ UTRs, 3’ UTRs, and coding sequence of transcripts using the largest isoforms detected in Pelechano et al. (2014).

**Figure 2–Figure supplement 2.**
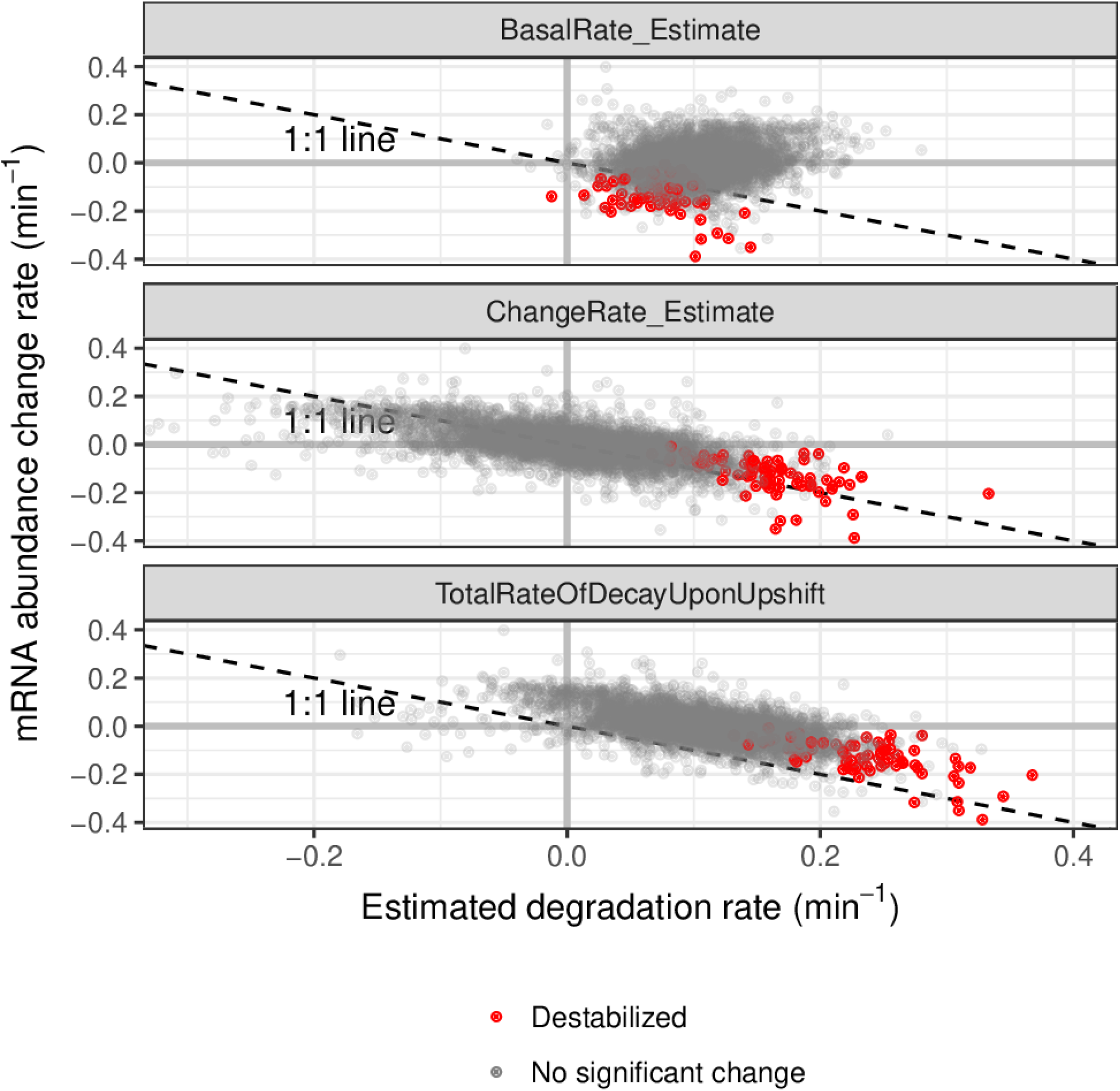
Comparisons of rates from this study with mRNA abundance change rates from Airoldi et al. (2016). Pre-upshift decay rates (top) don’t explain the abundance change. Decay rate refers to the rate of change, thus is the negative of the degradation rate. The degradation rate changes (difference between pre and post upshift) and the post-upshift rates (bottom) are anti-correlated with the abundance changes.

**Figure 2–Figure supplement 3.**
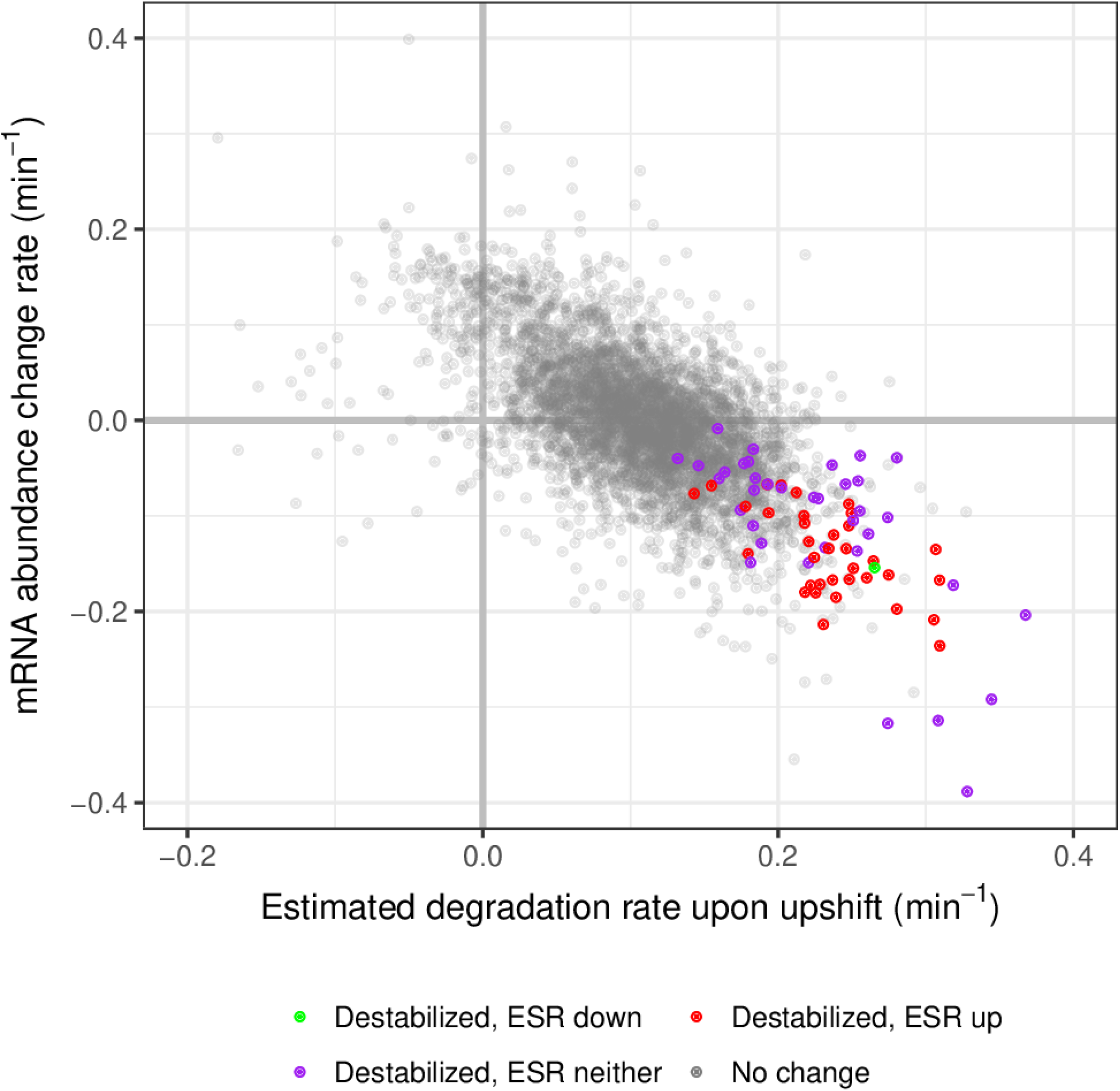
Comparisons of degradation rates from this study with mRNA abundance change rates from Airoldi et al. (2016). Destabilized transcripts are colored based on their membership in the ESR gene set, as described in the supplement of Brauer et al. (2008). Many of the destabilized set are ESR "up" genes, as they are increase in expression in response to stresses.

**Figure 2-Figure supplement 4.**
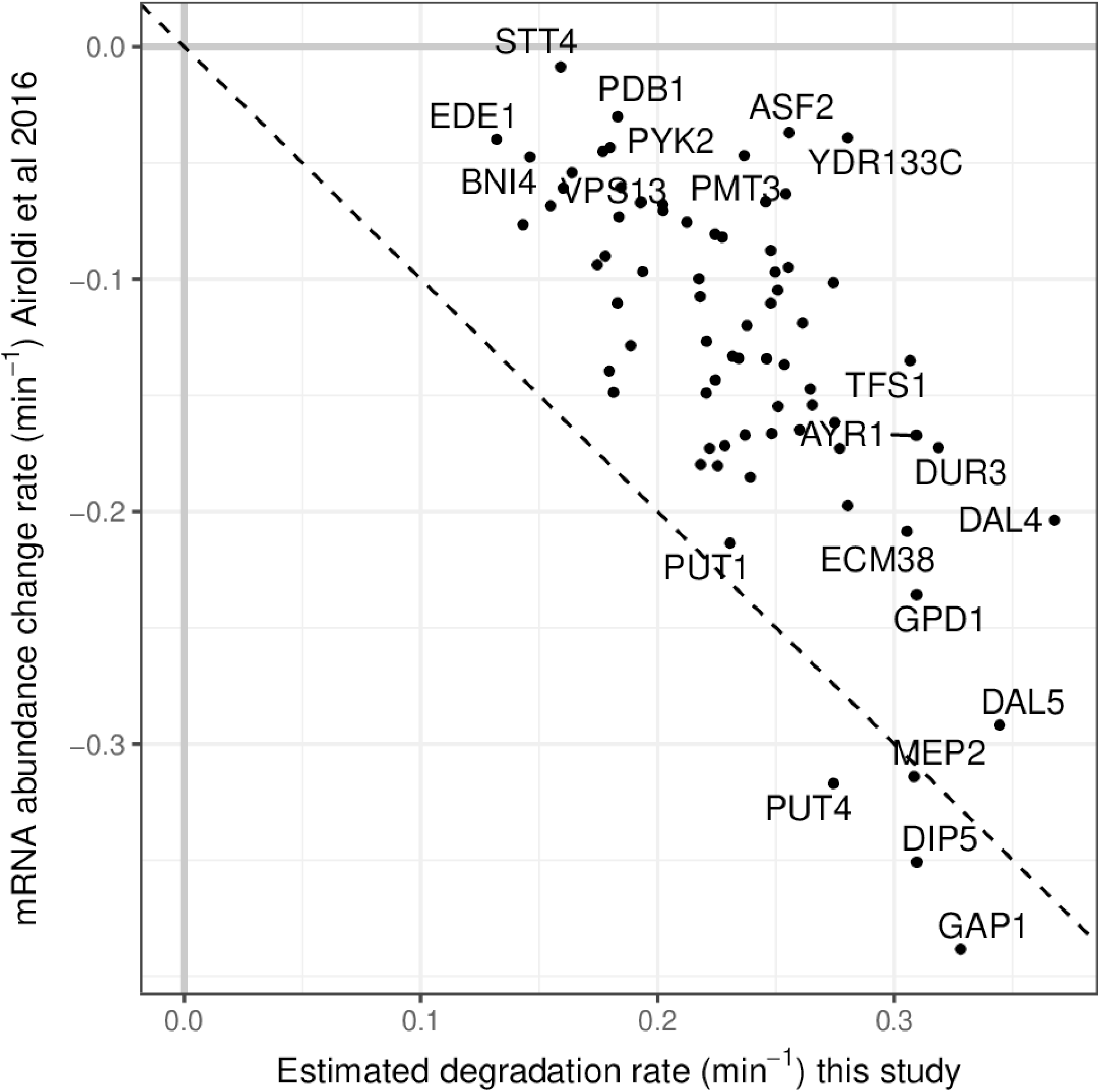
Scatter plot of significantly destabilized transcripts. For each, the x-axis is the fit rate of degradation rate post-upshift. On the y-axis is the mRNA abundance (expression) change rate (*Airoldi et al., 2016*) after the upshift. These values were modeled to normalized sequencing signal (x-axis) and normalized microarray ratio (y-axis). The dashed line is a 1:1 line of equality.

**Figure 2–Figure supplement 5.**
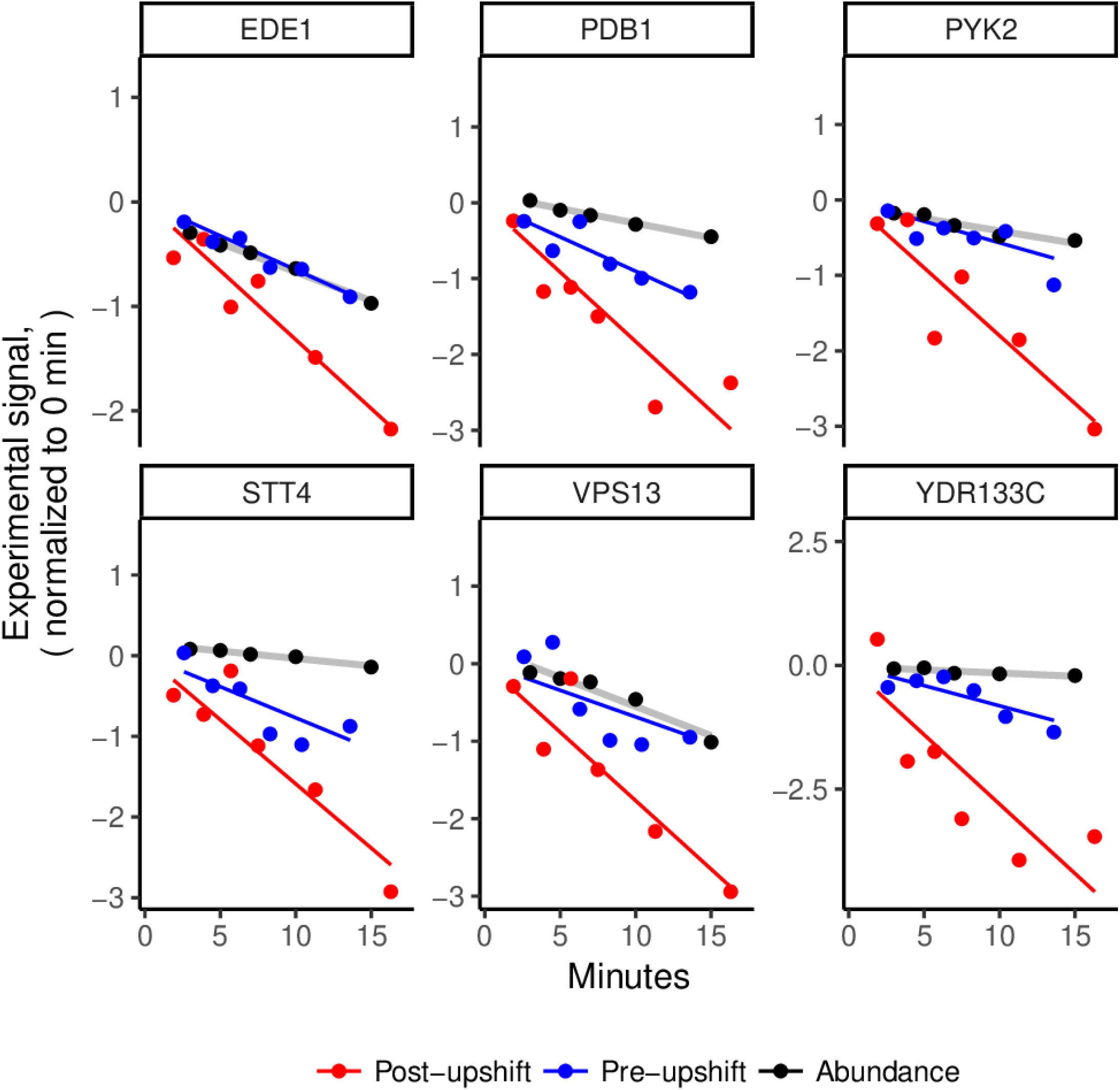
For several examples of the slowest decreasing (in the microarray fits) transcripts, we plot the microarray (abundance) and sequencing (decaying labeled abundance) data normalized to be on the same relative y-axis scale (subtracted t_0 y-intercepts of fits). Destabilization does not necessarily result in a rapid clearance of the mRNA.

**Figure 2–Figure supplement 6.**
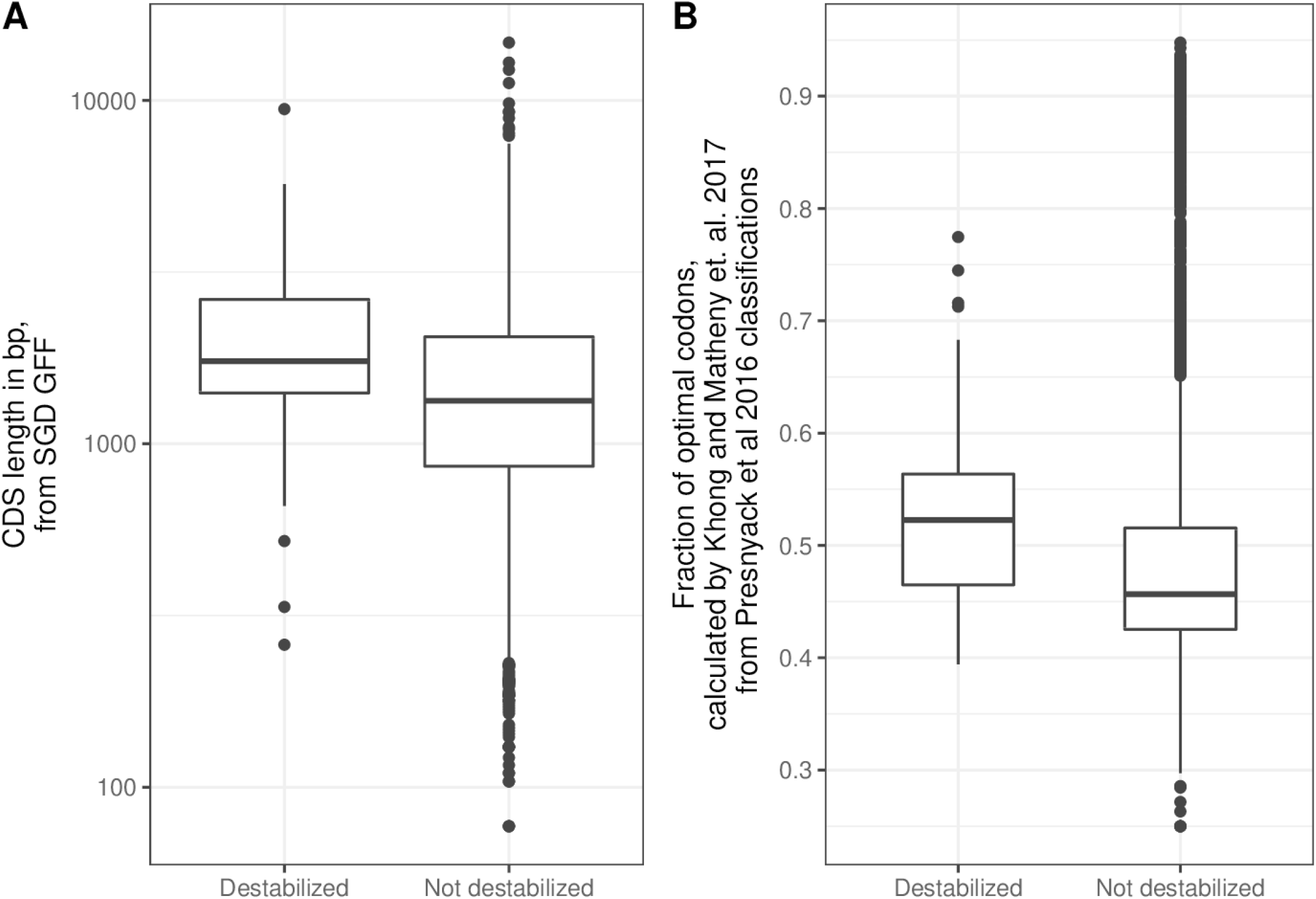
Comparisons of destabilized mRNAs with the rest of the transcriptome. a) Destabilized transcripts tend to have longer CDS lengths ( p-value < 2e-5 by Wilcoxon rank sum test). b) On average, the destabilized transcripts have more optimal codons than the rest of the transcriptome ( p-value < 2e-8 Wilcoxon rank sum test). The fraction of optimal codons per feature was obtained from the supplement of Khong et al. (2017) using definitions from Presnyak et al. (2015).

Figure2_Supplementary_Writeup.pdf

**Figure 2-Figure supplement 7.** Supplementary file with experimental rationale, details, and protocol for the label-chase experiment.

output/Figure2_Table_RawCountsTableForPulseChase.csv

**Figure 2-Figure supplement 8.** Raw counts of labeled mRNA quantified by RNAseq in label-chase experiment.

output/Figure2_Table_PulseChaseDataNormalizedDirectAndFiltered.csv

**Figure 2-Figure supplement 9.** Filtered label-chase RNAseq data for modeling, normalized directly within sample.

output/Figure2_Table_PulseChaseDataNormalizedByModel.csv

**Figure 2-Figure supplement 10.** Filtered label-chase RNAseq data for modeling, normalized by modeling across samples.

output/Figure2_Table_PulseChaseModelingResultTable_DirectNormalization.csv

**Figure 2-Figure supplement 11.** Degradation rate modeling results, from data normalized within samples.

output/Figure2_Table_PulseChaseModelingResultTable_ModelNormalization.csv

**Figure 2-Figure supplement 12.** Degradation rate modeling results, from data normalized across samples.

output/Figure2_Table_AcceleratedDegradationTranscripts_EnrichedGOandKEGGterms.csv

**Figure 2-Figure supplement 13.** Enriched GO and KEGG terms within the set of mRNA destabilized upon a nitrogen upshift, across sample normalization.

**Figure 3-Figure supplement 1.**
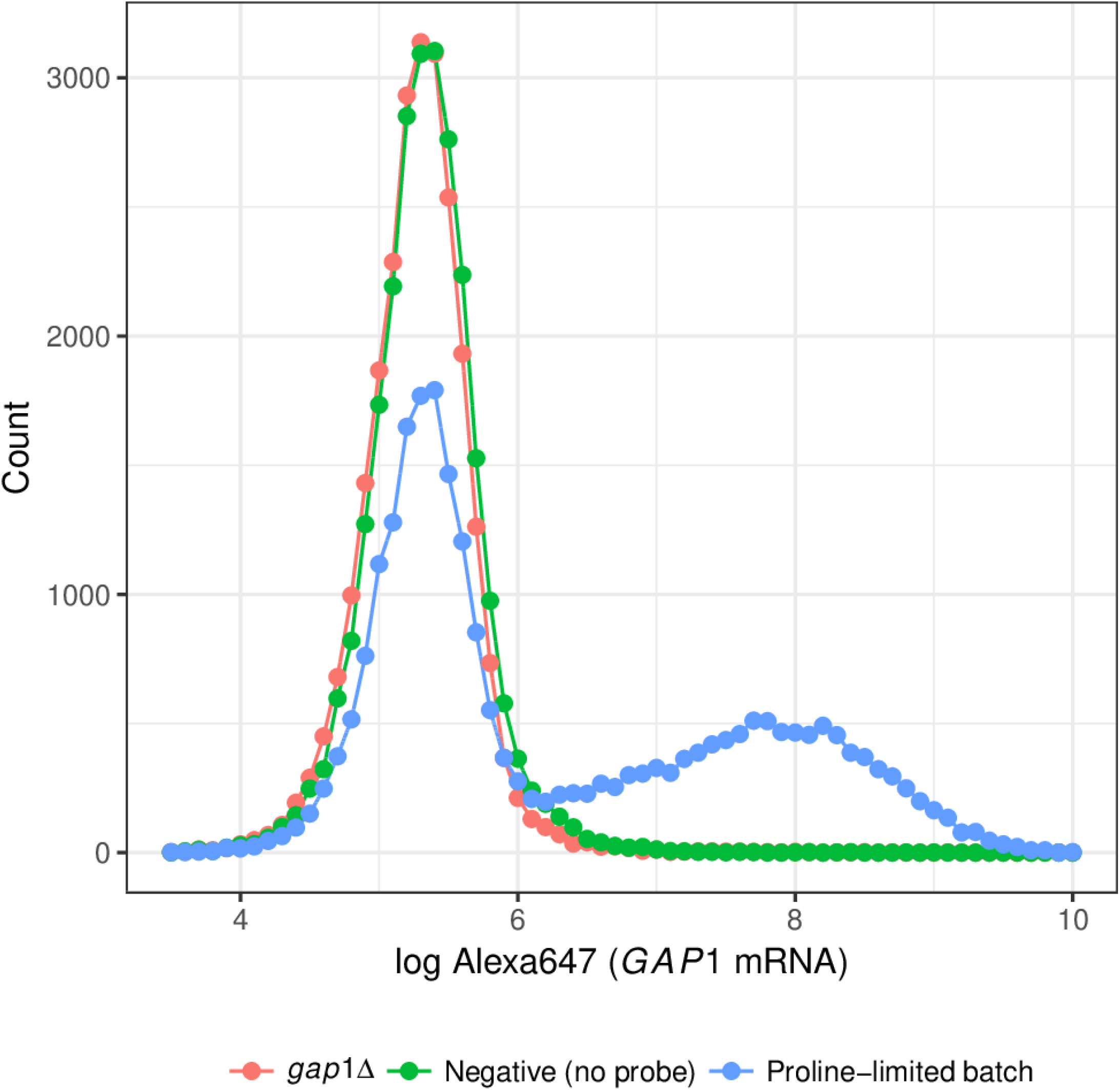
*GAP1* delete or omission of the targeting probe removes signal of GAP1 FISH. Wild-type or *GAP1*Δ cells were grown in proline-media, which induces expression of *GAP1.* As seen in the positive control, there is heterogeneity in the induction. This is likely due to technical issues, namely fixation time.

**Figure 4–Figure supplement 1.**
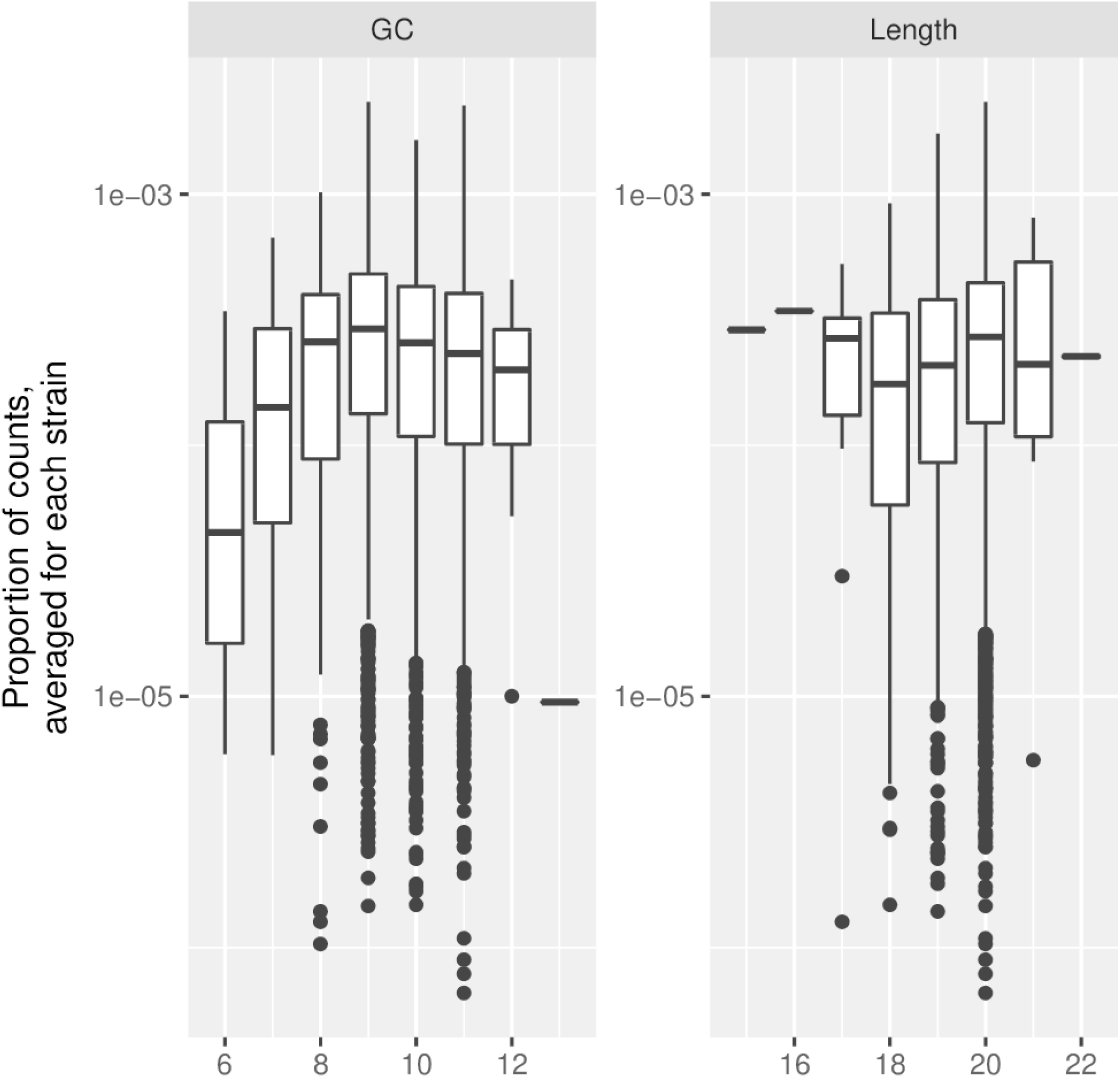
Distribution of proportion of counts by different counts of G or C, or by length. We see a complex relationship between relative counts and GC content, but no relation with length. We conclude that, as with any PCR-based sequencing assay, there exists a bias associated with their GC content.

**Figure 4–Figure supplement 2.**
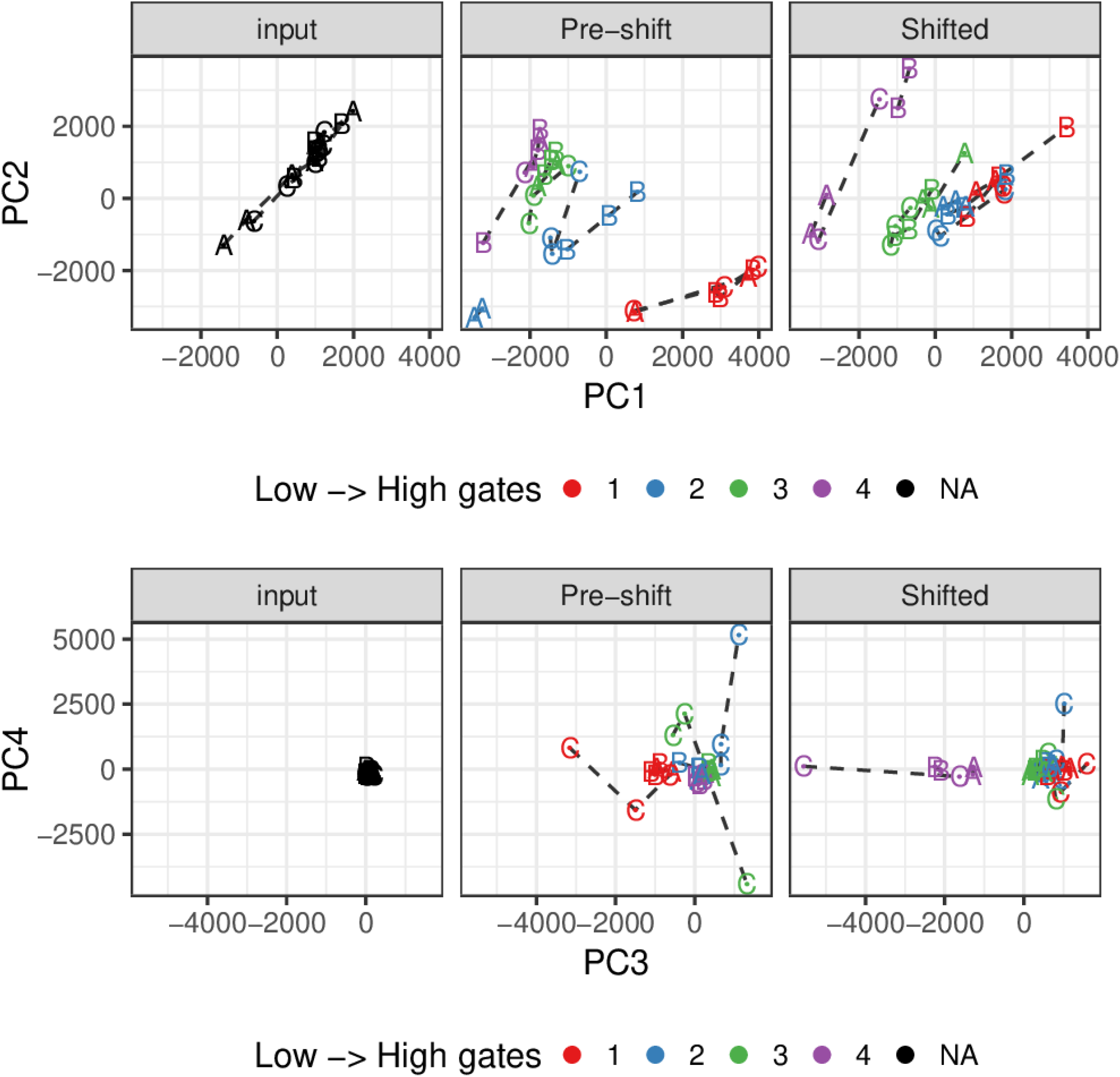
Principal components analysis of the abundance estimates for samples. Each color is a type of sample, from low to high gates (with black denoting the input samples before sort). Technical replicates are connected by dashed lines, biological replicates are each letter A B or C. At top, the first two prinicpal components show the separation of gates by signal intensity, and reflects that the lower gates on the upshifted samples were very close (blue and red samples on far right panel), within the distribution of the negative population. This is consistent with their tight sampling of the "GAP1-off" population, as seen in Figure 4a.

**Figure 4-Figure supplement 3.**
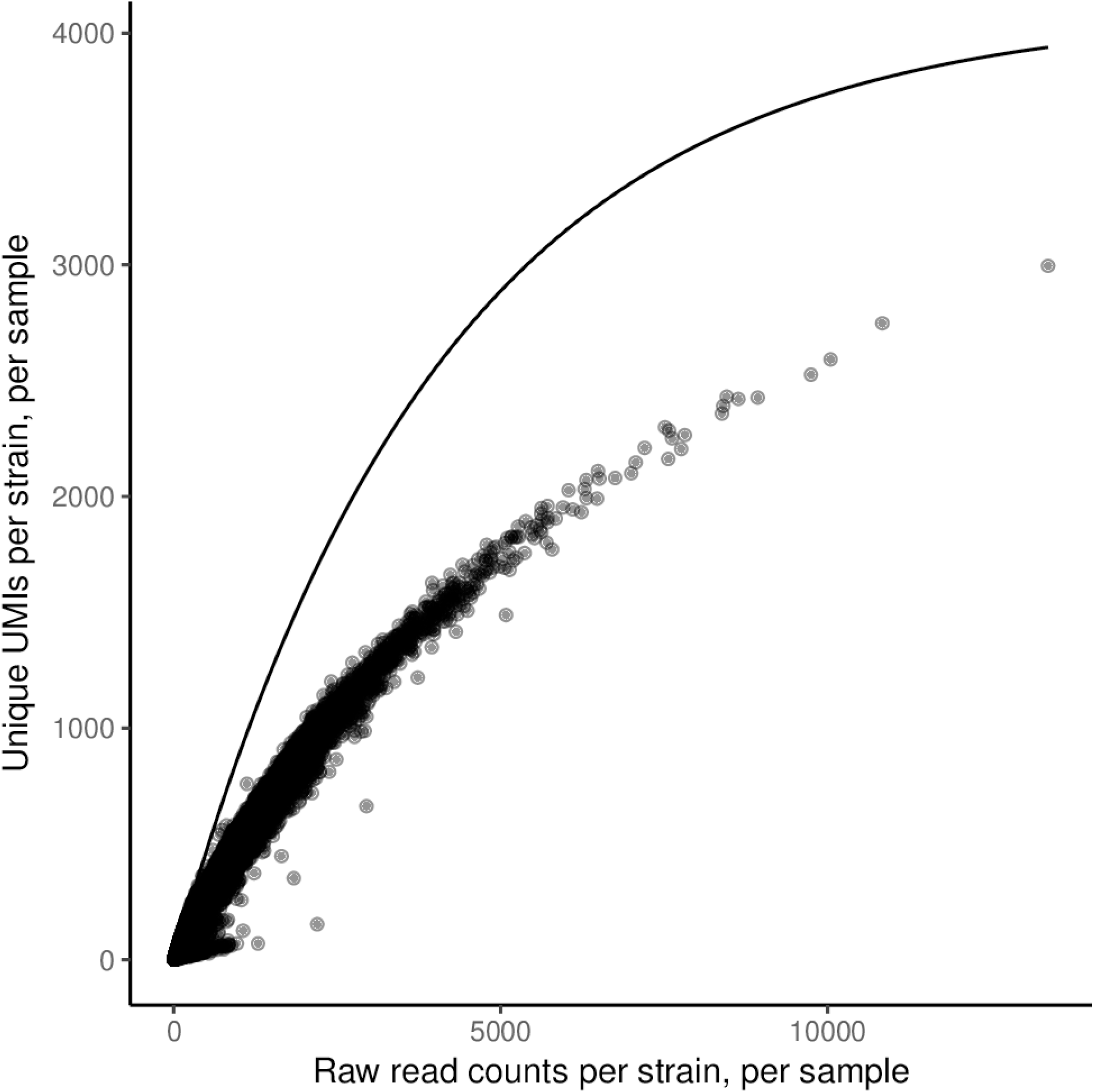
Rarefaction curve of UMI saturation. The solid-line curve denotes the theoretical expectation of total observations per UMI in a sample (x-axis) and the number of unique UMIs (y-axis). This curve shows how we expect UMI-collisions to depress the number of unique UMIs. Each point is from real data, with these two numbers tabulated for each combination of a sample and strain barcode. We see that these largely follow the curve of saturation of UMI-collisions, but that it falls well below the expectation of independent UMI-collision, thus we believe that there is an additional contribution of PCR-amplification noise (PCR duplicates).

**Figure 4–Figure supplement 4.**
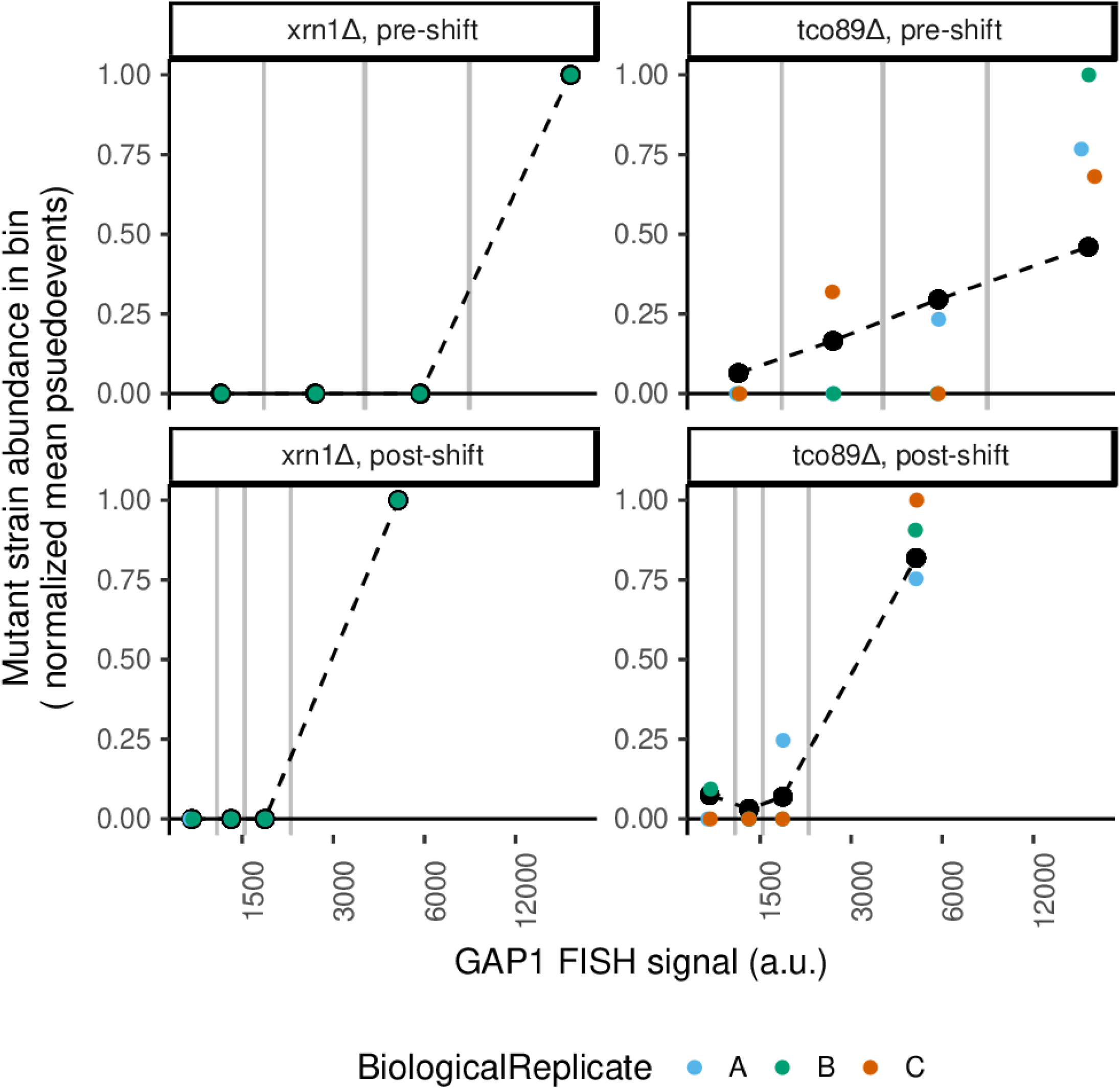
Data and fits for several mutants. xrn1Δ mutant (left) is lowly abundant in the library and is only observed in the highest bin of GAP1 signal, consistent with the role of Xrn1p as a global exonuclease. tco89Δ is the only detected member that would abrogate TORC1 activity. This mutant (right) has elevated GAP1 mRNA before and after the upshift, consistent with the role of TORC1 in repressing the NCR regulon.

Figure4_Supplementary_Writeup.pdf

Figure 4-Figure supplement 5. Supplementary file with experimental rationale, details, and protocol for the BFF experiment.

output/Figure4_Table_BFFcountsAndGateSettingsFACS.csv

Figure 4-Figure supplement 6. Raw counts of strain barcode quantification within each bin in the BFF experiment, and gate settings for the observations.

output/Figure4_Table_BFFmodelingData.csv

Figure 4-Figure supplement 7. BFF data filtered for modeling.

output/Figure4_Table_BFFallFitModels.csv

Figure 4-Figure supplement 8. The parameters of all models fit to the BFF data. output/Figure4_Table_BFFfilteredPooledModels.csv

Figure 4-Figure supplement 9. All 3230 models used for identifying strains with defective *GAP1* dynamics.

output/Figure4_Table_GSEanalysisOfBFFresults.csv

Figure 4-Figure supplement 10. Gene-set enrichment analysis results using *GAP1* estimates.

**Figure 5-Figure supplement 1.**
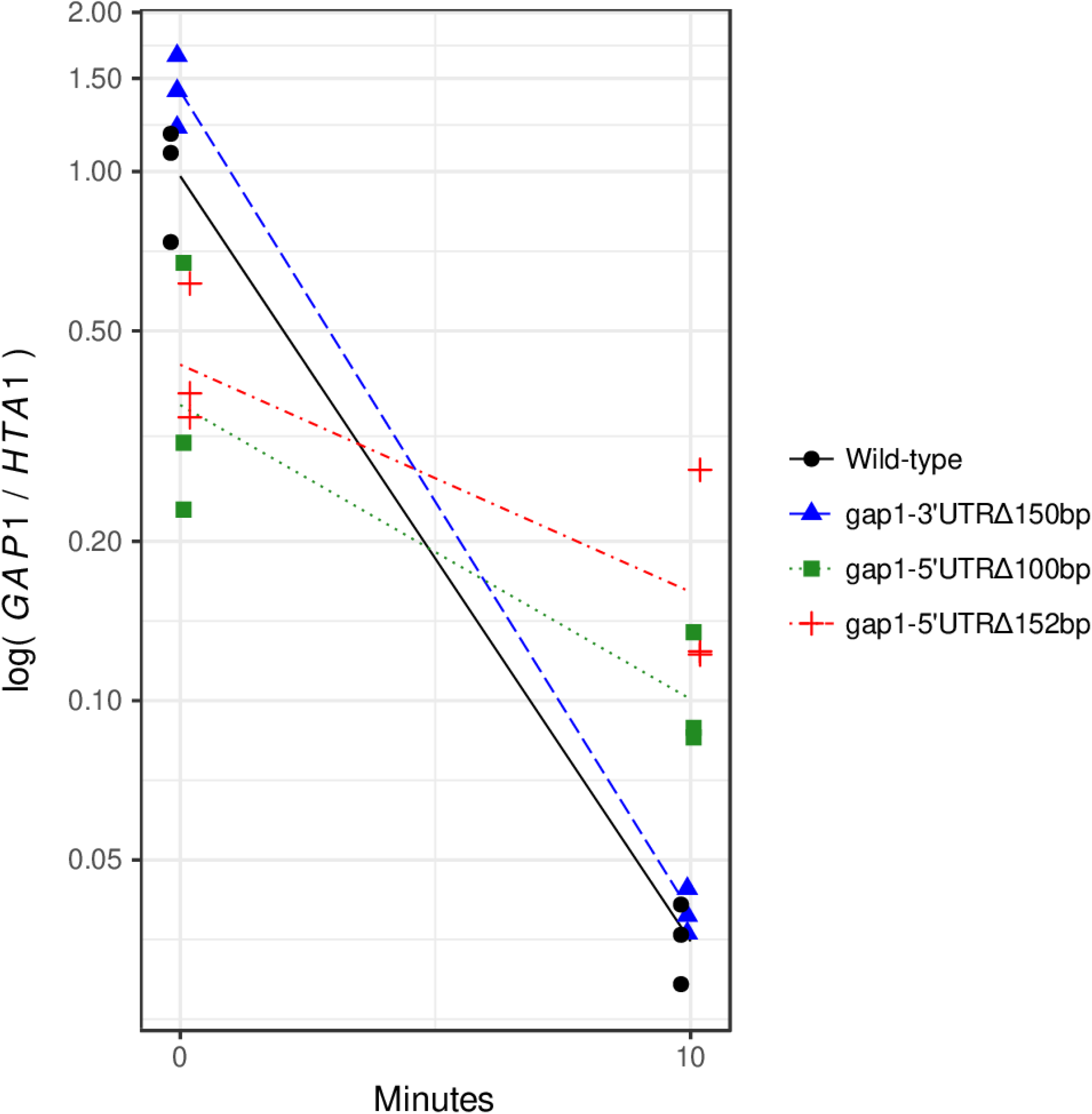
During strain construction, a deletion of 152bp 5’ of the start codon was also generated. We tested *GAP1* dynamics in this strain as well, and found that it shares the same phenotype as a 100bp deletion. Methods are the same as in Figure 5e, both 5’ deletes are slowed in clearance, ANCOVA p < 0.05.

**Figure 5-Figure supplement 2.**
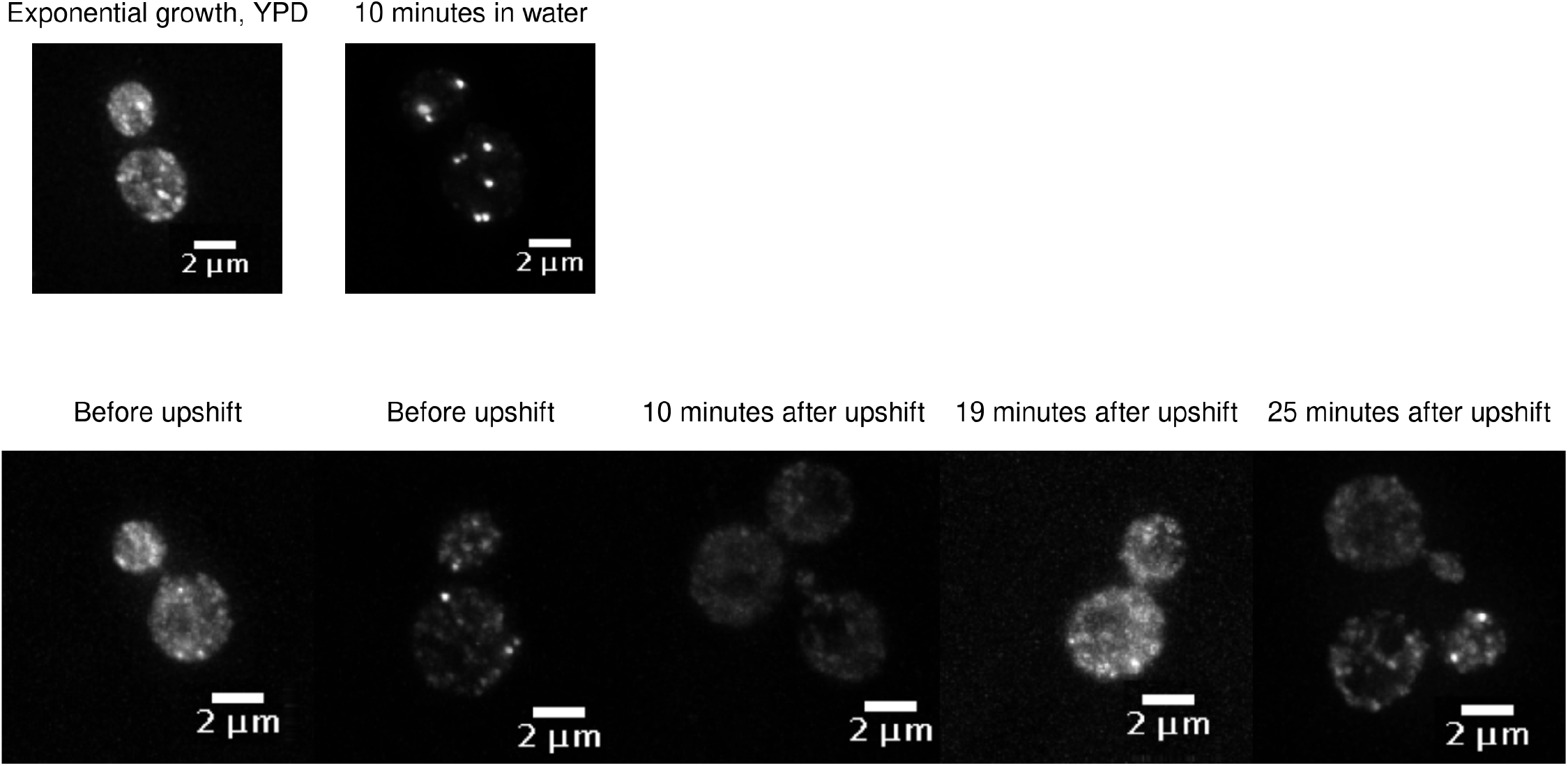
A strain harboring a copy of Dcp2p-GFP expressed from a plasmid was grown in conditions of exponential phase in YPD or 10 minutes of starvation in water (first row). This is a common condition known to result in the formation of processing-body foci of Dcp2-GFP. We do not see either formation or dissolution of Dcp2-GFP foci during the nitrogen upshift.

**Figure 5–Figure supplement 3.**
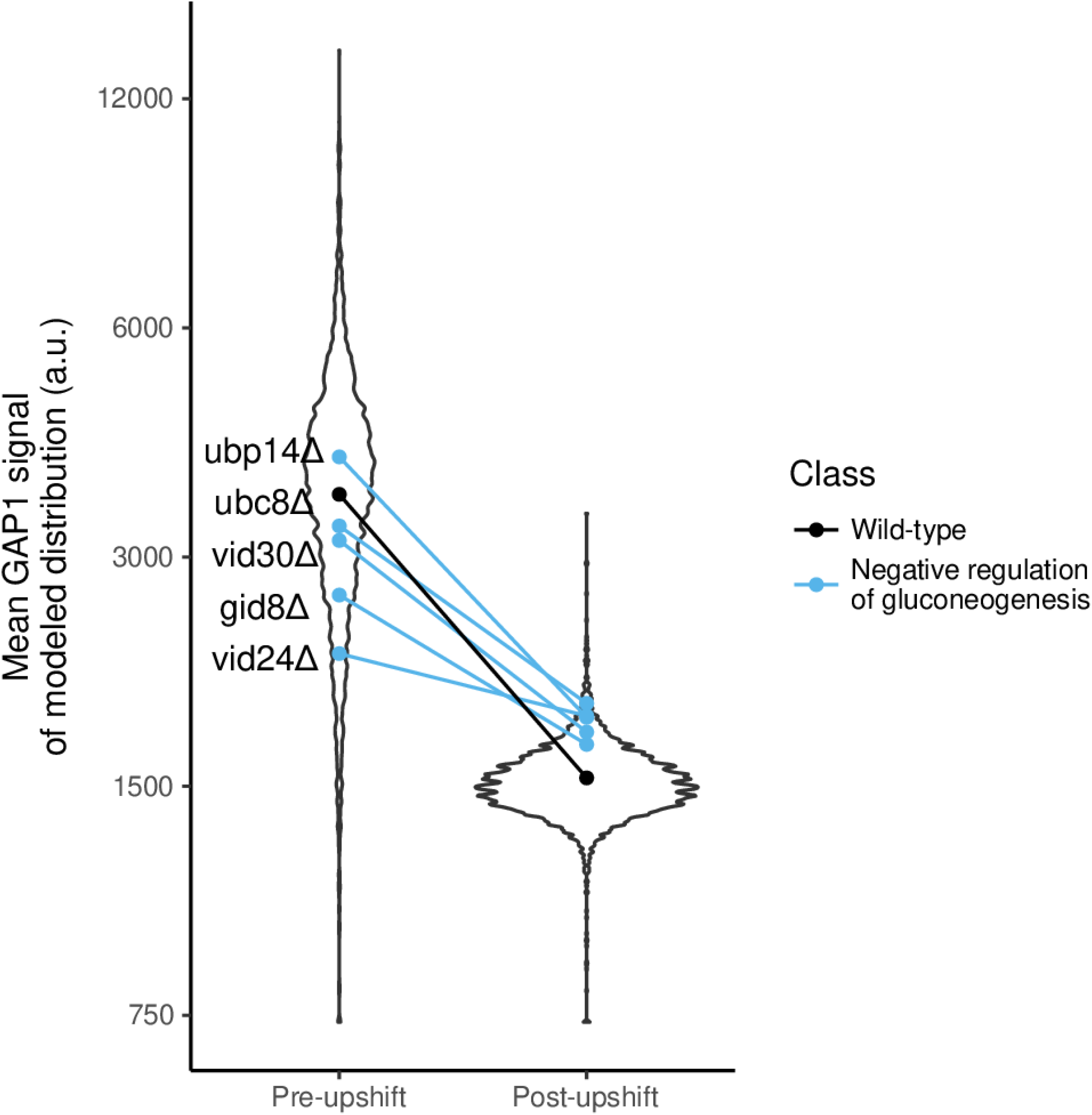
Knock-out mutants of negative regulators of gluconeogenesis are associated with higher estimated GAP1 mean after the upshift, by GSEA analysis of GO-terms (p-value < 0.05).

**Figure 5–Figure supplement 4.**
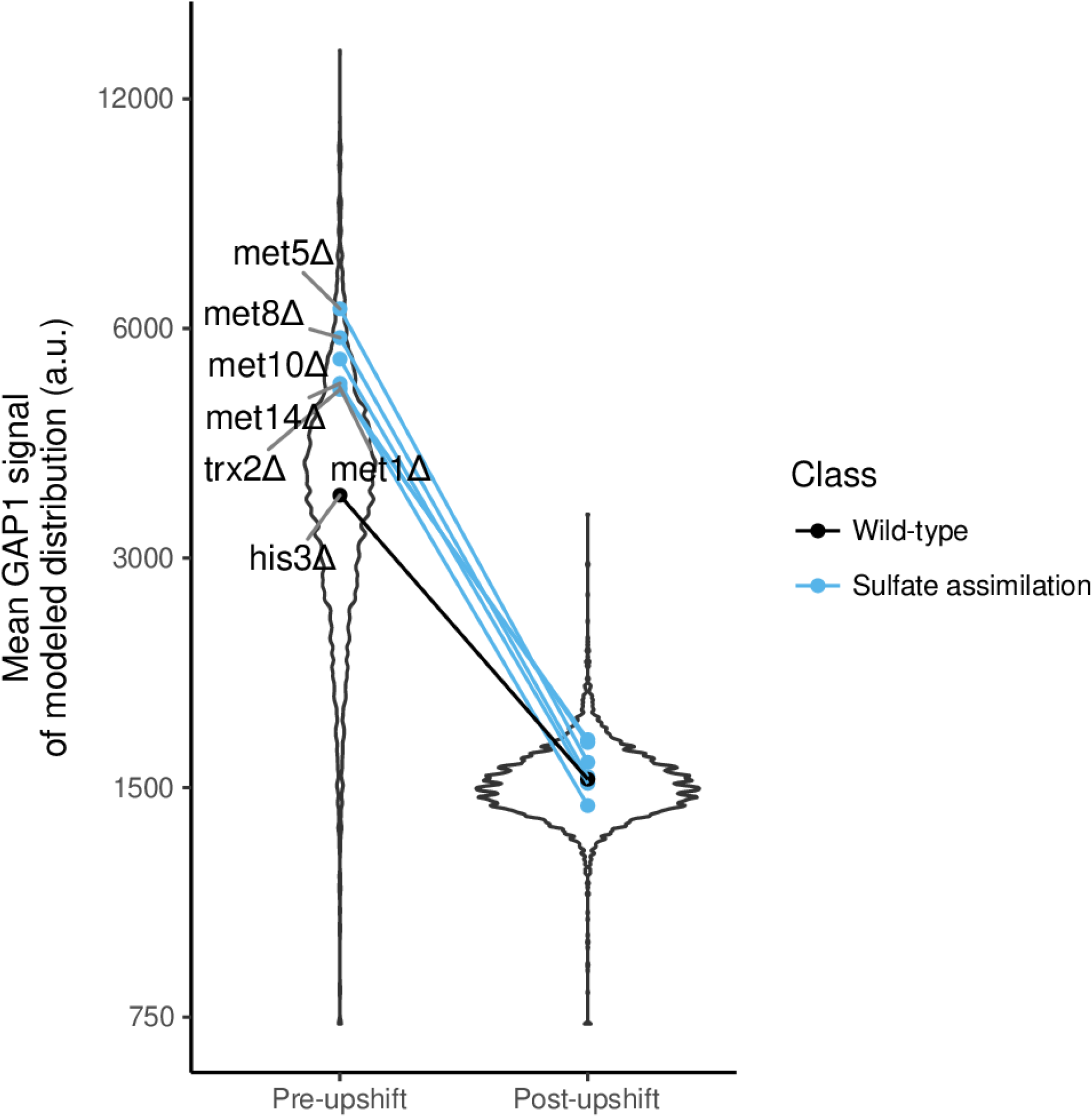
Knock-out mutants of involved in sulfate assimilation are associated with higher estimated GAP1 mean before the upshift, by GSEA analysis of GO-terms (p-value < 0.05).

**Figure 5–Figure supplement 5.**
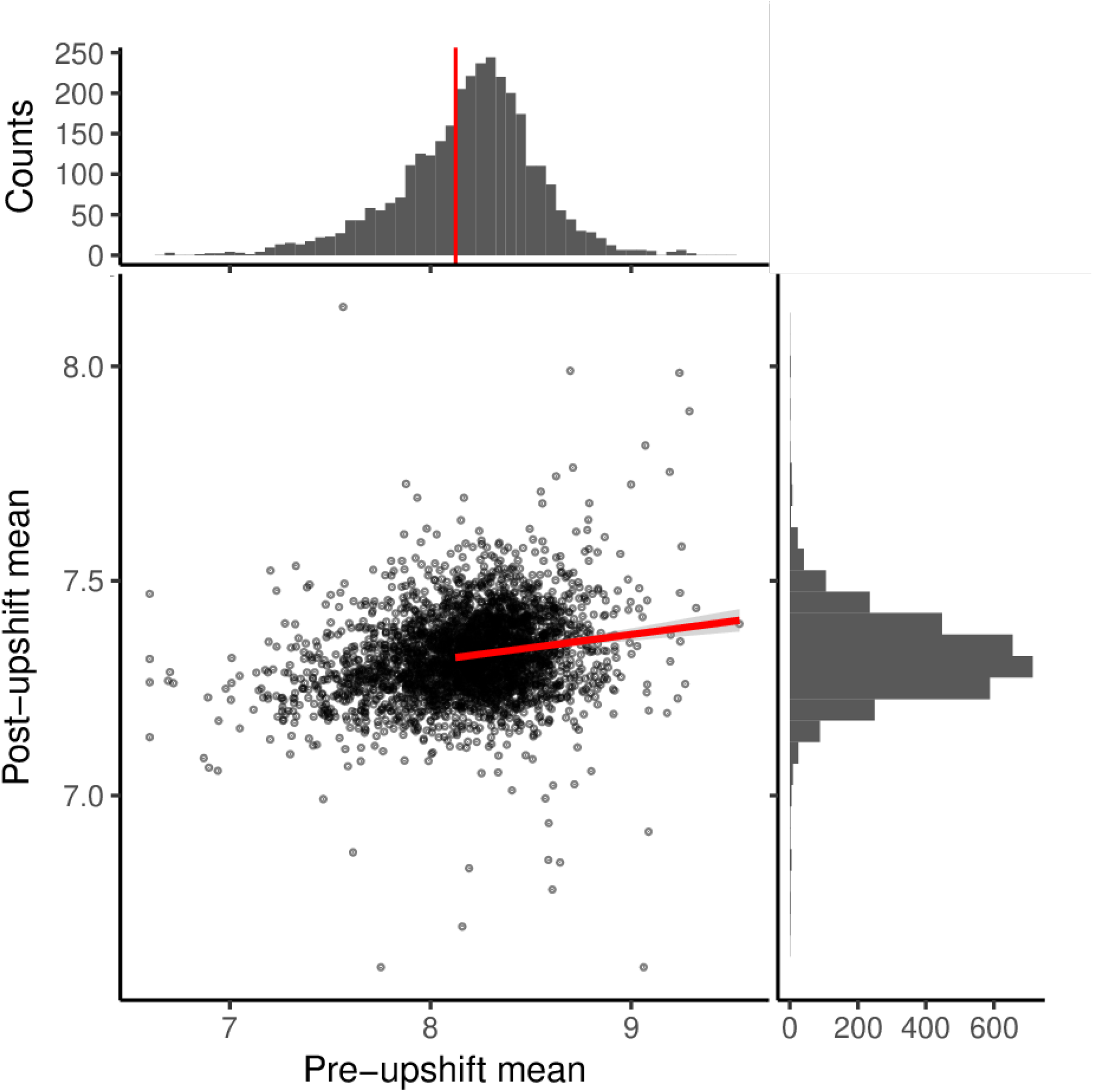
The relationship between the estimated mean before the upshift and after the upshift. Scatter plot of the estimated means, with marginal histograms along top and right. Red vertical line on top histogram is a cut-off of GAP1 mRNA induction for this analysis, and is the mean of the fit to wild-type minus the standard deviation of that distribution. The red linear regression line is fit to all points above this threshold, in which expression was detected before the upshift.

